# Reference genome bias in light of species-specific chromosomal reorganization and translocations

**DOI:** 10.1101/2024.06.28.599671

**Authors:** Marius F. Maurstad, Siv Nam Khang Hoff, José Cerca, Mark Ravinet, Ian Bradbury, Kjetill S. Jakobsen, Kim Præbel, Sissel Jentoft

**Affiliations:** Centre for Ecological and Evolutionary Synthesis, Department of Biosciences, University of Oslo, Oslo, Norway; Fisheries and Oceans Canada, Newfoundland, St John’s, Canada; Norwegian College of Fishery Science, The Arctic University of Norway, Tromsø, Norway

## Abstract

Whole-genome sequencing efforts has during the past decade unveiled the central role of genomic rearrangements—such as chromosomal inversions—in evolutionary processes, including local adaptation in a wide range of taxa. However, employment of reference genomes from distantly or even closely related species for mapping and the subsequent variant calling, can lead to errors and/or biases in the datasets generated for downstream analyses. Here, we capitalize on the recently generated chromosome-anchored genome assemblies for Arctic cod (*Arctogadus glacialis*), polar cod (*Boreogadus saida*), and Atlantic cod (*Gadus morhua*) to evaluate the extent and consequences of reference bias on population sequencing datasets (approx. 15-20x coverage) for both Arctic cod and polar cod. Our findings demonstrate that the choice of reference genome impacts population genetic statistics, including individual mapping depth, heterozygosity levels, and cross-species comparisons of nucleotide diversity (π) and genetic divergence (D_XY_). Further, it became evident that using a more distantly related reference genome can lead to inaccurate detection and characterization of chromosomal inversions, i.e., in terms of size (length) and location (position), due to inter-chromosomal reorganizations between species. Additionally, we observe that several of the detected species-specific inversions were split into multiple genomic regions when mapped towards a heterospecific reference. Inaccurate identification of chromosomal rearrangements as well as biased population genetic measures could potentially lead to erroneous interpretation of species-specific genomic diversity, impede the resolution of local adaptation, and thus, impact predictions of their genomic potential to respond to climatic and other environmental perturbations.

## Introduction

Recent advancement within sequencing technologies and bioinformatic tools have revolutionized the field of biology. Pioneering studies have been conducted within human genomics, which have improved our understanding of biological processes tremendously. The number of studies on wildlife and marine species is also increasing^1–4^, and over the past years, several larger international initiatives have been established to characterize all of life’s genomic diversity^5^. Within these efforts, the overall goal is to generate highly contiguous reference genomes (i.e., chromosome level) that can be used in a i) comparative setting to describe the genomic diversity between species, and/or conduct ii) within-species genome-wide characterization of cryptic ecotypes and sub-population differentiations^4–7^.

While the number of high-quality reference genomes is growing, there is still a shortage in the number of reference genomes available for various taxa^8^. In the cases where a reference genome for the focal species is missing, the standard method is to select a close relative for mapping and subsequent variant calling^9^. When using a reference genome from a distantly related species (or a divergent population), the genomic divergence between the reference and the target species can impact mapping, variant calling, and downstream inferences^9–14^. For instance, measures of heterozygosity—important measures for conservation genomics—can be overestimated when employing more divergent references^10–13^. However, few studies have examined how discrepancies in genomic architecture between the reference and target species would impact the identification of e.g., larger structural variants, such as chromosomal inversions. Since the beginning of the genomics area, chromosomal inversions have been recognized as part of the standing genomic variation of a species, and/or sub-populations/ecotypes, that are likely to play important roles in evolutionary processes, including local adaptation^15–20^. For instance, in Atlantic cod (*Gadus morhua*; L., 1758), four larger chromosomal inversions are found to discriminate between populations throughout its geographical distribution, i.e., dominating the observed genomic divergence by large allele frequency shifts^15,21–23^. It is suggested that these are of high importance for maintaining the genomic divergence between locally adapted populations as well as the iconic migratory Northeast Arctic cod (NEAC) and the more stationary Norwegian coastal cod (NCC)^15,21–24^. Would such and other structural variants be overlooked or inadequately characterized due to larger or smaller inter-chromosomal reorganizations between the reference used and the focal species? In an earlier study conducted on European plaice (*Pleuronectes platessa*), a difference in number of putative chromosomal inversions were recorded based on using the species-specific reference vs. using the Japanese flounder (*Paralichthys olivaceus*)^25,26^ that potentially could be due to species-specific differences in number of inversions and/or other types of inter-chromosomal reorganizations.

Within the gadids, major genomic reorganizations and reshufflings have been documented, and especially within the two cold-water specialists: the Arctic cod (*Arctogadus glacialis;* Peters, 1872) and the polar cod (*Boreogadus saida;* Lepechin, 1774)^27^. Additionally, for polar cod a large number of polymorphic chromosomal inversions (with the potential impact on sub-population structuring) have been detected^28^. Such major genomic reorganizations and reshufflings could potentially lead to downstream bioinformatic errors in mapping, variant calling, and data interpretation, depending on the selection of reference. In this study, we aimed at taking the full advantage of the newly generated chromosome-anchored genome assemblies of the closely related Arctic cod^27^, polar cod^27^, and NEAC^29^ to assess how the selected reference genome impacts the mapping depth, heterozygosity level and measures of population differentiation and divergence between Arctic cod and polar cod, when exploring population-level data of the two species collected from the northern Barents Sea and adjacent regions (Figure 1b). Additionally, we investigated how the different reference genomes influence the detection of chromosomal inversions, focusing exclusively on the Arctic cod. Both Arctic cod and polar cod represent important sympatric species inhabiting the Arctic, one of the world’s most rapidly changing environments that is undergoing warming at a pace almost four times faster than the global average^30^. Until now, there are only a few studies that have looked into the population genetic structuring of Arctic cod and polar cod using a handful of genetic markers^31–36^ and even fewer that have used whole genome sequencing approaches^28,37^, and by such, this study will advance our insight into the genomic composition and potential within these species in the light of the ongoing climatic changes.

**Figure 1:**
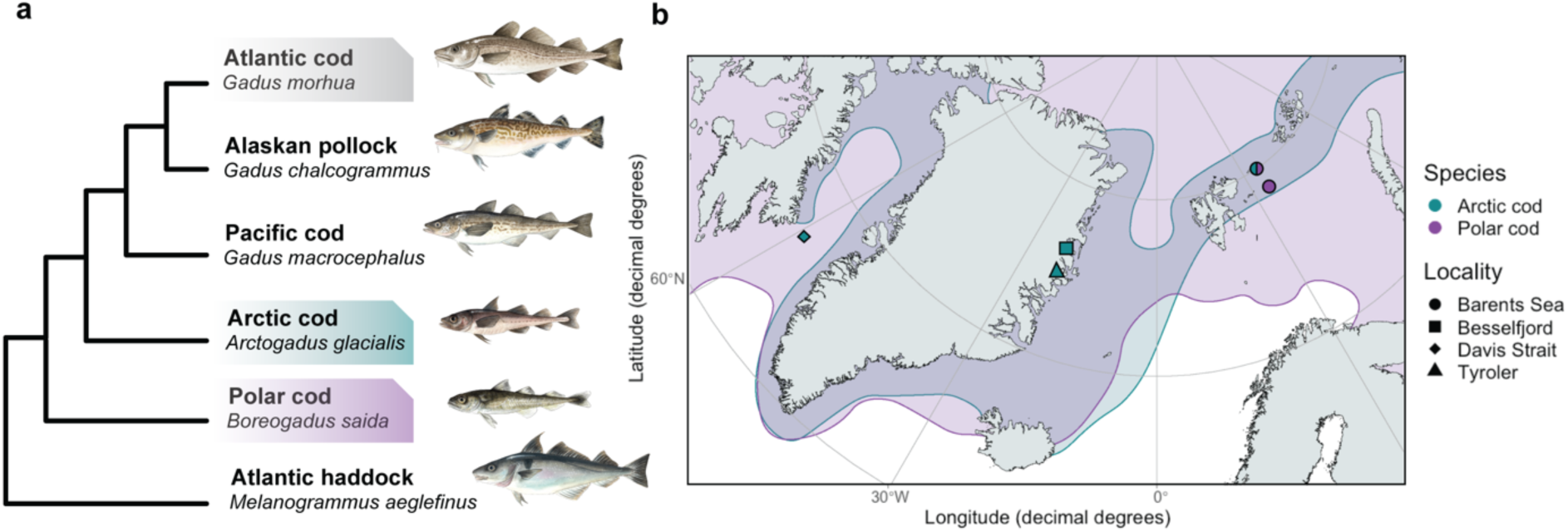
Phylogenetic relationship and distribution of Arctic cod and polar cod. a) Phylogenetic relationship of Arctic cod redrawn from Matschiner et al.^24^ and Hoff et al.^27^ The phylogenetic placement of Arctic cod is not fully resolved, as it may be either a sister lineage to *Gadus* or a sister species to polar cod^24,27^. Species used as reference genomes in this study are highlighted. b) Map of sampling localities of Arctic cod and polar cod with their distributions in the sampling region redrawn from Mecklenburg et al.^69^ Illustrations by Alexandra Viertler.

## Materials and Methods

### Sample acquisition and sequencing

The collection of Arctic cod (N=14, Table S1) used in this study was obtained via the TUNU- cruises (UiT, The Arctic University of Norway) and from other international collaborators, including N=11 individuals from Northeast Greenland (Tyroler and Besselfjord) and N=2 individuals from Canada (Davis Strait), as well as one specimen collected in the Barents Sea (Figure 1b). The collection of polar cod (N=14, Table S1) is a subset from a larger dataset^28^ from the northern Barents Sea (Figure 1b). DNA isolation for Arctic cod was done by following the QIAGEN DNeasy Blood & Tissue kit protocol. DNA concentration measurement, library preparation, and sequencing were performed by the Norwegian Sequencing Centre. See Supplementary Sequencing Report for more information.

### Study design

The whole genome sequencing data were used to generate three *cross-species* datasets where data from both Arctic cod (N=14) and polar cod (N=14) were included (Figure 2a), as well as three *intraspecific* datasets where we focused on the Arctic cod samples (Figure 2b). Both sample collections (i.e., *cross-species* and *intraspecific*) were mapped against the reference genomes of either i) Arctic cod^27^, ii) polar cod^27^, and iii) Northeast Atlantic cod (NEAC)^29^, with the main purpose to assess the choice of reference on mapping depth as well as heterozygosity levels. Additionally, population genetic measures, such as nucleotide diversity (π), genetic differentiation (F_ST_), and genetic divergence (D_XY_) were estimated to assess the influence of reference genome choice in a *cross-species* context. Moreover, we utilized the *intraspecific* datasets to assess the precision in detection of chromosomal inversions within Arctic cod. This was conducted by comparing the degree of overlap between the inversions detected when using either the Arctic cod (i.e. the benchmark) vs. the polar cod or the NEAC genome as a reference.

**Figure 2:**
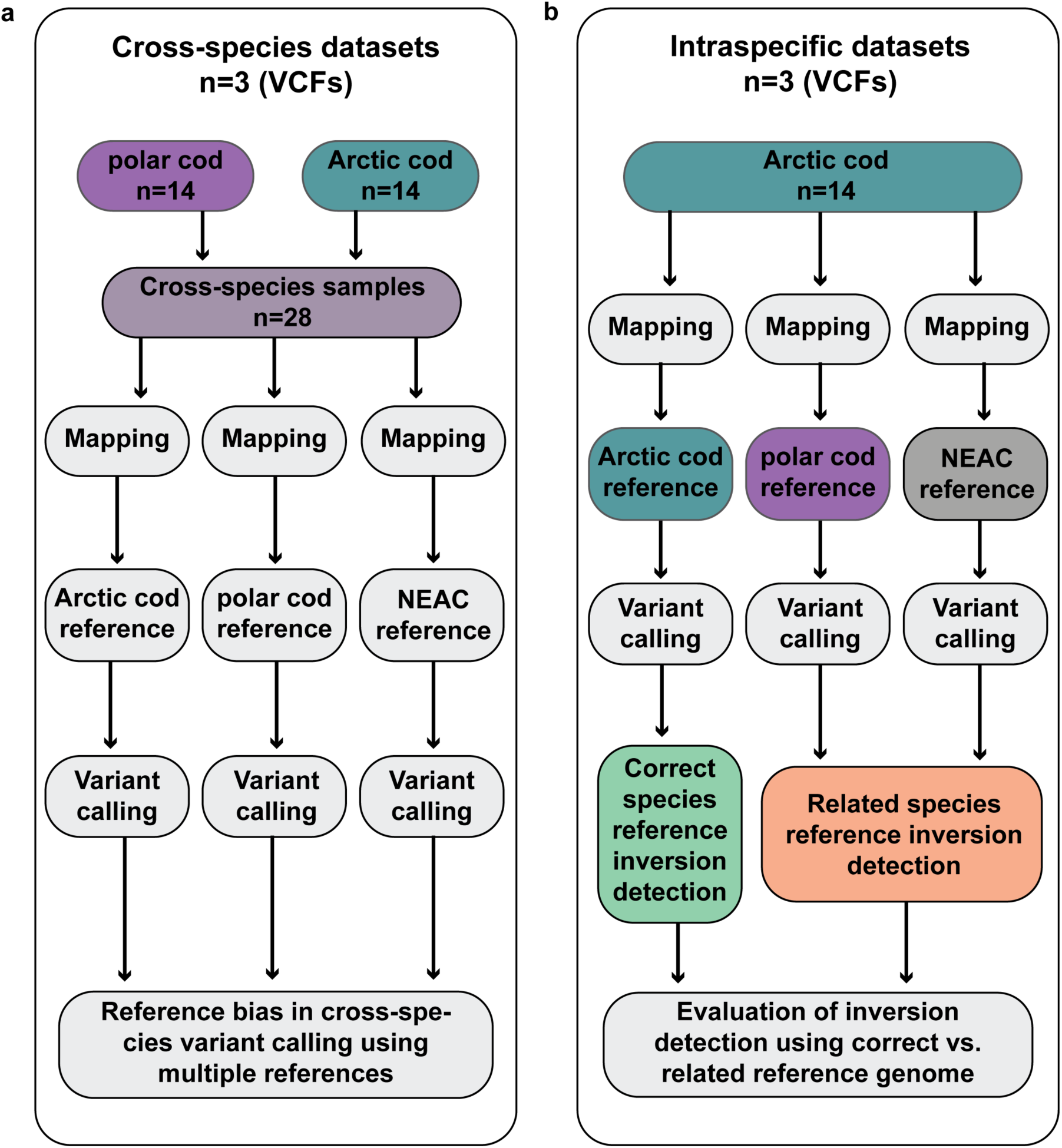
Flowchart of the sample design and generation of the *cross-species* and *intraspecific* datasets. a) For the generation of the three *cross-species* VCFs, we used samples of Arctic cod (N=14) and polar cod (N=14). Each sample was individually mapped against three different reference genomes: Arctic cod, polar cod, and (Northeast Arctic cod) NEAC. After mapping, the samples were grouped based on the reference genome they were mapped against. This approach was employed to assess the extent of reference bias in *cross-species* variant calling when using different reference genomes. b) To investigate the impact of reference bias on inversion detection, we generated three *intraspecific* datasets focusing on Arctic cod samples (N=14). These samples were mapped against the same three reference genomes used for the *cross-species* VCFs. In this analysis, the Arctic cod reference was considered the accurate benchmark for detecting inversions. The detected inversions were then compared to those identified when using a related species’ reference genome, to evaluate the influence of reference choice on inversion detection.

### Mapping and variant calling

To obtain the six separate datasets (i.e., three VCFs for the *cross-species* analysis and three VCFs focusing on the *intraspecific* analysis) we started by trimming Illumina PE reads using Trimmomatic v0.39^38^ with default settings. Mapping to the different references was done using the Burrows-Wheeler Alignment Tool v0.7.17^39^ (BWA-MEM algorithm) with default settings. Alignment files for each sample were merged and sorted using SAMtools v1.9^40^. Duplicated reads were marked using MarkDuplicates v2.22.1^41^. Variant calling was performed using the Genome Analysis Toolkit (GATK) v4.2.0.0^42^. For this, each mapped sample was individually called into GVCFs using HaplotypeCaller. GVCFs for individual samples were then combined into the six different VCFs, as described above in the experimental design, and imported into a GenomicsDataBase using GenomicsDBImport. Joint genotyping was performed using the GenotypeGVCFs tool to produce final VCFs. Single nucleotide polymorphisms (SNPs) were extracted and downsampled to 100,000 SNPs using SelectVariants to make diagnostic plots for filter parameter evaluation. Filtering was done by following the GATK hard-filtering recommendations and manually inspecting the diagnostic plots as suggested in https://speciationgenomics.github.io/. After the initial round of filtering, we used VCFtools v0.1.16^43^ to retain only biallelic sites (see Table S2 and S3 for filtering parameters and Table S4 for the number of SNPs after filtering). Lastly, in-depth inspection of the datasets generated was conducted using PLINK v1.9^44^ and VCFtools v0.1.16. for detection of potential data biases (for more information see Supplementary Note 1). A summary of the workflow is shown in Figure 2.

### Evaluation of population structure, mapping, and variant calling based on reference used

We analyzed read depth distributions of mapped reads for Arctic cod and polar cod samples against the three references using mosdepth v0.2.4^45^ in fast mode, with a window size of 500 bp. Additionally, VCFtools v0.1.16 was used to evaluate the proportion of heterozygous sites per sample. The population genetic structure between and within the two species was investigated using PLINK v1.9 to perform a Principal Component Analysis (PCA), using both the *cross-species* and the *intraspecific* datasets.

For an evaluation of the genetic diversity detected within the *intraspecific* datasets, we also carried out demographic inference and estimated female effective population size (N_e_) for Arctic cod using BEAST v2.6.7^46^ under the Bayesian skyline model^47^. The analysis was done twice, once only with Arctic cod samples in the present study (N=14) and including Arctic cod (N=33) samples sourced from NCBI (see Supplementary Note 2 for more details). Additionally, for the *cross-species* datasets, π, F_ST_ ^48^, and D_XY_ between Arctic cod and polar cod were estimated using pixy v1.2.6^49^, applying a window size of 10,000 bp.

### Detection of chromosomal inversions in Arctic cod

For the *intraspecific datasets* (i.e., the three intraspecific VCFs mapped to the three different reference genomes) detection of chromosomal inversions was performed using complementary approaches. The workflow is illustrated in Supplementary Figure 5. First, we used a PCA-based approach following Huang et al^17^. This involved quantifying genetic variation within each chromosome using the R package lostruct in windows of 50 SNPs^50^. When conducting PCAs of inversions, heterokaryotypes are expected to cluster between the two homokaryotype clusters for individuals carrying alternative inversion orientations^51^. Thus, resulting lostruct plots were manually checked for regions along chromosomes where the PCA for the MDS corners displayed three distinct clusters. After detecting potential inversion regions, VCFtools v0.1.16 was used to extract the regions harboring the inversion signal and calculate the heterozygosity for each sample. PLINK v1.9 was then used to calculate a new PCA of the SNPs within this region. In the cases where the PCA displayed an inversion signal, clusters were assigned to either homokaryotypes with most individuals (common group), heterokaryotypes as the group clustering in the center (het group), or homokaryotypes with the fewest individuals (rare group). Due to the low sample count for Arctic cod, the heterozygosity distribution could not be plotted using conventional boxplots, instead we used a binning strategy implemented in the ggplot2 function geom_dotplot^52^.

Next, F_ST_ and D_XY_ were calculated using pixy v1.2.6 in windows of 10,000 bp between the rare and common groups along chromosomes to assess patterns of genetic differentiation and divergence outside and inside potential inversion regions. Lastly, patterns of linkage disequilibrium (LD) were investigated for the chromosomes that displayed potential signals of inversions. The expectation for chromosomes harboring inversions is that regions within the inversion will show high LD among all samples (when both homokaryotypes are present) but not among samples with the same inversion orientation^17^. As the calculation of LD in a pairwise fashion for whole chromosomes produces millions of data points, SNPs had to be down-sampled. PLINK v1.9 was used to remove sites with more than 0.01% missing data, and SNPs were randomly thinned down to 10% of the original count. After thinning of SNPs, PLINK v1.9 was used to calculate LD in a pairwise fashion for the SNPs left within the chromosome of interest. Due to the high number of data points still left, the R package scattermore was used to produce the LD plots^53^. We used the MDS plots along the chromosomes to define the boundaries of the inversions and corroborated with the LD patterns.

### Synteny between the three references

To investigate chromosomal rearrangements synteny analysis between the Arctic cod, polar cod, and NEAC references was done using a syntenic block analysis with McScanX^54^. The result of the synteny analysis was visualized on the Synvisio interactive homepage^55^.

## Results & Discussions

### Genetic structure of Arctic cod and Arctic cod vs. polar cod

The PCA conducted on the *intraspecific* genomic dataset revealed a separation among the Arctic cod specimens along the first principal component (PC1) axis, explaining 10.7-10.9% of the variation in the datasets depending on the reference used (Figure 3a-c). Additionally, a separation along the PC2 axis was demonstrated, explaining 8.31-8.42% of the variation in the datasets (Figure 3a-c). When inspecting this separation against the various variant calling statistics (Figure S1-S3), we found that neither mean depth nor presence of missing sites appeared to have a notable influence on the positioning of the samples within the PCA. Mean depth was generally consistent across most samples, except for a single individual from Davis Strait. This individual, sourced from a publicly available dataset (Table S1), had been sequenced to a greater depth (approx. 30x coverage) than the others. Furthermore, among the samples, one individual from Besselfjord displayed a higher degree of missing data compared to the rest. The proportion of heterozygous sites, however, tended to overlap to some degree with the sample positioning along the PC1 axis. It should be noted that this was not the case for all samples, for instance, the Davis Strait sample (with the highest coverage and highest proportion of heterozygotic sites present) was placed in the middle of the gradient (Figure S1-S3). Additionally, the proportion of heterozygous sites was generally higher using either polar cod or NEAC vs. Arctic cod as reference but did not impact the placement of the samples within the PCA (Figure 3a-c; Figure S1-S3). It is therefore tempting to speculate that a sub-population structuring within Arctic cod is present. However, to fully assess this and define the different sub-populations a larger dataset with more individuals from a larger geographical range is needed.

**Figure 3:**
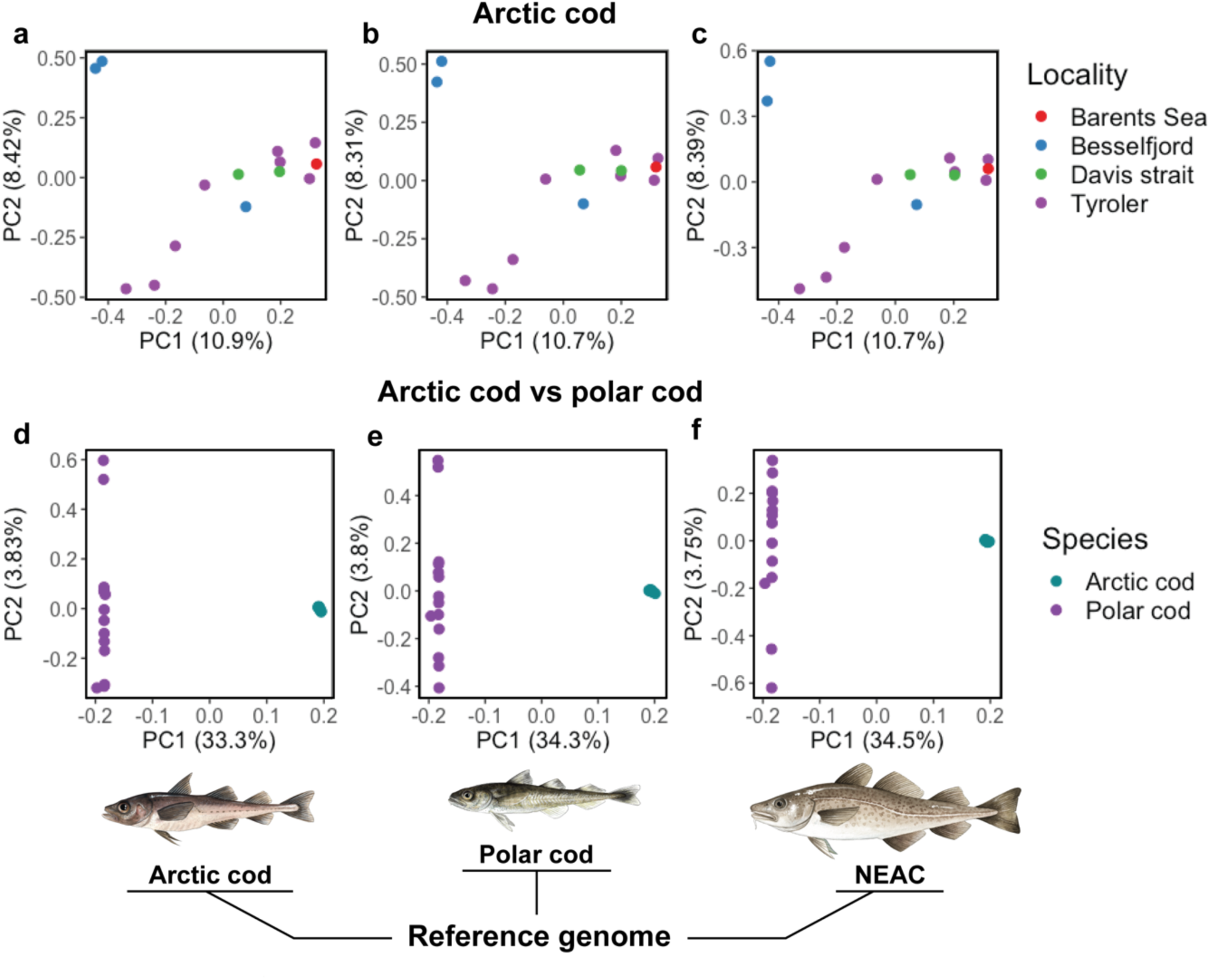
Genetic structure for Arctic cod samples (*intraspecific*) and Arctic cod vs. polar cod (*cross-species)* using the three different reference genomes. The map shows the different sampling localities of Arctic cod and polar cod used in this study. a, b, c) PCA of Arctic cod samples against the three references, and d, e, f) PCA of *cross-species* datasets using the three references. The three references Arctic cod, polar cod, and NEAC used for the PCAs shown below.

The PCAs on the *cross-species* datasets uncovered a distinct clustering pattern irrespective of the reference used, where the samples clustered in accordance with their respective species (Figure 3d-f), i.e., one cluster for Arctic cod and one cluster for polar cod, respectively. Additionally, a difference in how the two species clustered along the PC2 axis was detected, with Arctic cod exhibiting minimal intraspecific variation, whereas polar cod displayed intraspecific variability along the PC2 axis (Figure 3d-f), explained by 3.75-3.81% of the variation in the datasets, depending on the reference used. Taken together, these findings indicate that polar cod has a larger standing genetic variation compared to Arctic cod, which could be linked to the difference in female N_e_ observed between the species (Figure S4 and Hoff et al.^27^) as well as documented by others^37,56,57^.

### Impact of reference genome on mapping and variant calling statistics

For the *cross-species* datasets, the estimation of mean mapping depth uncovered a species-specific variability, which was dependent on the reference genome used. We detected highest mean depth when individual sequencing data were mapped against their intraspecific reference, while using one of the two other codfishes as the reference resulted in lower mean depth (Figure 4a). Lowest mapping depth was observed using NEAC as the reference, i.e., the most distant reference with lowest sequence identity and thus, the lowest potential mappability for both the Arctic cod and polar cod datasets. Notably, the polar cod datasets displayed higher overall depth levels, irrespective of reference used, due to the fact that the polar cod samples were sequenced in a separate batch with slightly higher coverage (see Supplementary Materials and Methods in Hoff et al.^28^). Moreover, the proportion of heterozygous sites estimated (Figure 4b) mirrored the patterns of mean depth observed, where the lowest number of heterozygous sites was detected when the intraspecific reference was employed (Figure 4b), while a higher proportion of heterozygous sites was detected when one of the two heterospecific codfishes was used as the reference (Figure 4b). Thus, a higher mean depth resulted in a lower proportion of heterozygote sites and vice versa. Intriguingly, the degree of heterozygosity, seemed to be less impacted when using the more distantly related NEAC as a reference. Even if having the lowest mapping depth, the heterozygosity level was not as pronounced as seen when using either Arctic cod or polar cod as the reference (Figure 4a and b). These findings could potentially be coupled to the high genomic content of short tandem repeats detected within codfishes^58–60^ combined with the GadMor3 genome assembly being of higher quality and more contiguous compared to the Arctic cod and the polar cod genome assemblies^27,29^. Mapping towards these lower quality genomes would potentially result in a higher degree of erroneous mapping of reads, i.e., misalignments (especially within the repetitive regions) vs. when mapping towards the higher quality NEAC genome assembly. Accordingly, the lower quality of the Arctic cod and the polar cod genomes, i.e., with a lower resolution of the repetitive regions, combined with higher sequence identity between these two species, could easily result in higher mapping depth (as documented above), as well as a higher degree of wrongly called heterozygous sites^10–14,61^. It should also be noted, that our findings could be explained by the fact that NEAC is genomically more divergent vs. the two other species, resulting in lower mappability and lower number of callable sites, and thus, less heterozygote sites detected. But, based on the similar number of sites called using the different references (see Table S4), the latter explanation seems less plausible.

**Figure 4:**
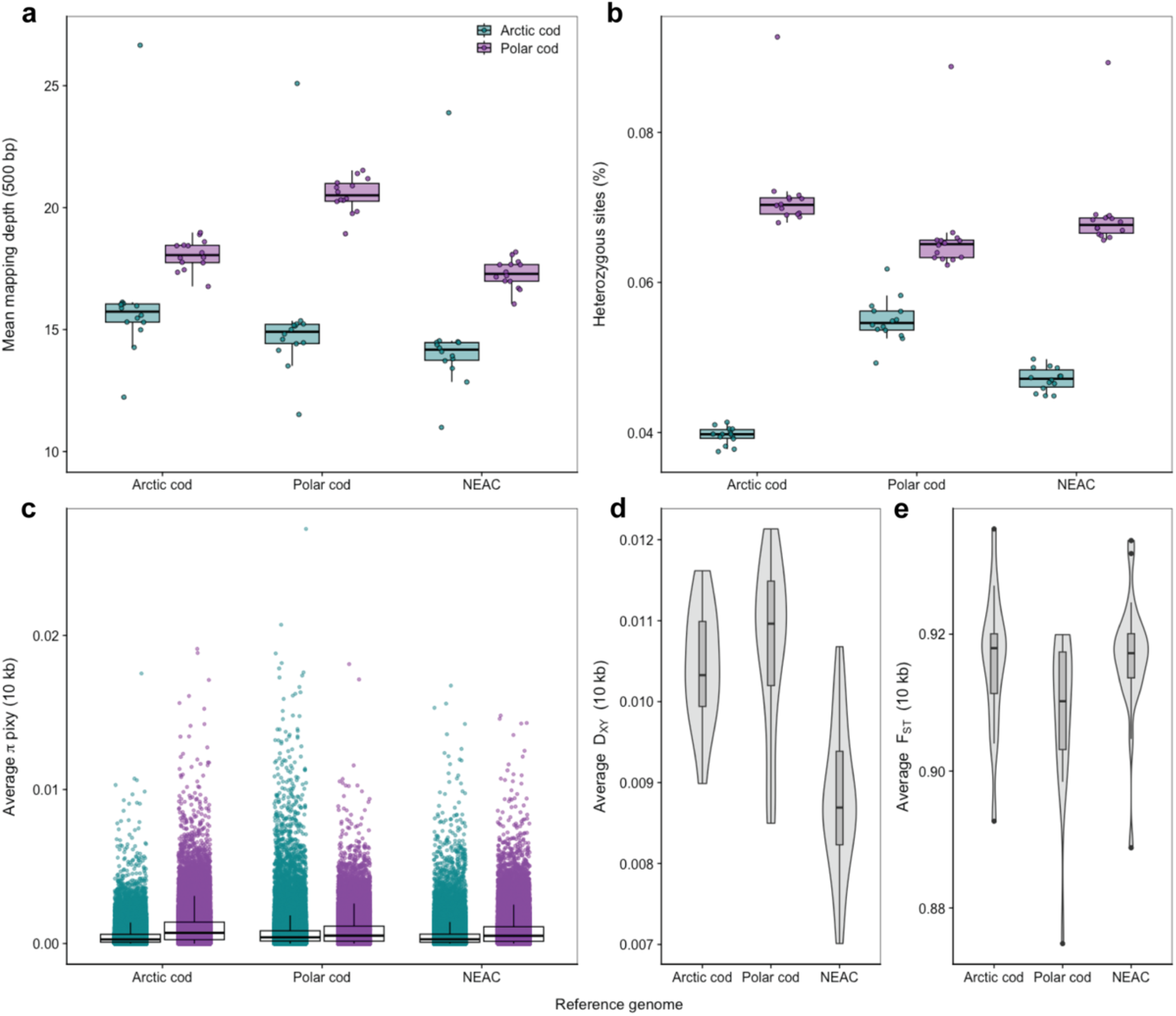
Variability in sample statistics and population measures for the cross-species comparison using different references for Arctic cod and polar cod. a) The mean mapping depth differs for Arctic cod and polar cod samples based on the reference chosen, i.e., Arctic cod, polar cod or NEAC. The highest mean depth is seen in samples when they are mapped against their respective intraspecific references. b) Similarly, the proportion of heterozygous sites per sample, calculated using VCFtools after variant calling, also changes with the reference used. The lowest values are found in Arctic cod and polar cod when analyzed against their own intraspecific references. c) Average π values in windows across the three different reference genomes for each species, calculated using pixy, demonstrate variation in calculated π values depending on the reference used. d, f) Average D_XY_ and F_ST_ for each chromosome in the cross-species comparison of Arctic cod and polar cod, calculated using pixy, using the three different references, also show variability depending on the reference chosen.

For the population genetic statistics calculated for each of the species, we discovered varying results depending on the reference used (Figure 4c-e). The average π estimates displayed similar overall trends regardless of the reference genome used (Figure 4c; box plots). However, when either Arctic cod or polar cod was used as a reference, the non-reference species in the *cross-species* datasets exhibited a tailing of the average π values (Figure 4c; points). In contrast, using NEAC as a reference, the tailing appeared less pronounced and more similar to the estimates seen for the intraspecific comparisons. Similarly, the average background D_XY_ divergence (Figure 4d) between the species was higher when Arctic cod and polar cod were used as references, while a notable decrease in genetic divergence was observed when employing NEAC as the reference. These observations combined, could probably also be linked to the difference in quality of the genome assemblies, with the NEAC having the highest quality and lower degree of misalignments and/or due to poorer mappability, as discussed above. Additionally, the employment of an equally distant relative as reference for both species, could here be an asset, i.e., not introducing any reference bias towards one of the species when performing the variant calling. Such a bias could most likely influence the genetic diversity detected between the two species, seemingly resulting in an overestimation of the genetic divergence between Arctic cod and polar cod, when compared to the results achieved when using NEAC as the reference. On the other hand, when using NEAC as the reference there might be a higher chance that the polymorphic sites and the divergence detected between the two species are located within conserved regions (where the mappability is better), which could lead to an underestimation, as observed in our comparisons (Figure 4d). Contradictory to π and D_XY_, calculation of average background F_ST_ differentiation between Arctic cod and polar cod uncovered a similarly high degree of fixation between the species, irrespective of which of the three references used (Figure 4e). The rather large interspecific differentiation at the whole genome-wide level corroborates the findings from the PCA analyses (Figure 3d, e and f), indicating that the reference used does not impact the variant calling to any degree to determine the global degree of differentiation between the species, when using F_ST_ and/or PCA analyses. In contrast, genetic diversity and genetic divergence, measured by π and D_XY_, are seemingly more sensitive to the choice of reference used.

### Detection of multiple chromosomal inversions in Arctic cod

For the *intraspecific* dataset when using Arctic cod as reference genome we detected six chromosomal inversions that fulfilled the criteria defined by our inversion detection protocol (Figure S5). The inversions detected were found on chromosome 1, 6, 10, 11, 13, and 14, spanning from 2 Mb to 14 Mb in size (Table 1; Figure 5; Figure S6-S10). Furthermore, we detected five additional putative inversions, i.e., regions displaying the same patterns as the other inversions, but with weaker LD signals, less clear heterozygosity distribution, and/or only 2 or less individuals in the rare cluster (Table 1; Figure S11-S17). Among the putative inversions, the ones identified on chromosome 7 represented a special case where two smaller regions in the center of the chromosome exhibited inversion signals but did not share the same individuals between clusters (Figure S11 and S12), and thus, denoted as two separate putative inversions. The absence of an inversion signal in the intermediate region further supports two independent inversions (Figure S13). On chromosome 9, we detected a signal indicating a putative inversion. However, this region did not display distinct R2 values along the LD heatmap (Figure S14). Lastly, putative inversions were detected on chromosomes 3 and 15, respectively, were both fulfilled all steps for inversion detection but only had a single sample in the rare cluster (Figure S15 and S16). Additionally, for chromosome 10, we identified a region upstream of the inversion that also displayed a high degree of differentiation (Figure S17). However, this upstream region lacked the distinct PCA clusters and typical heterozygosity distribution expected for inversions (Figure S17), and therefore, was not classified as part of this inversion nor as a separate putative inversion.

**Figure 5:**
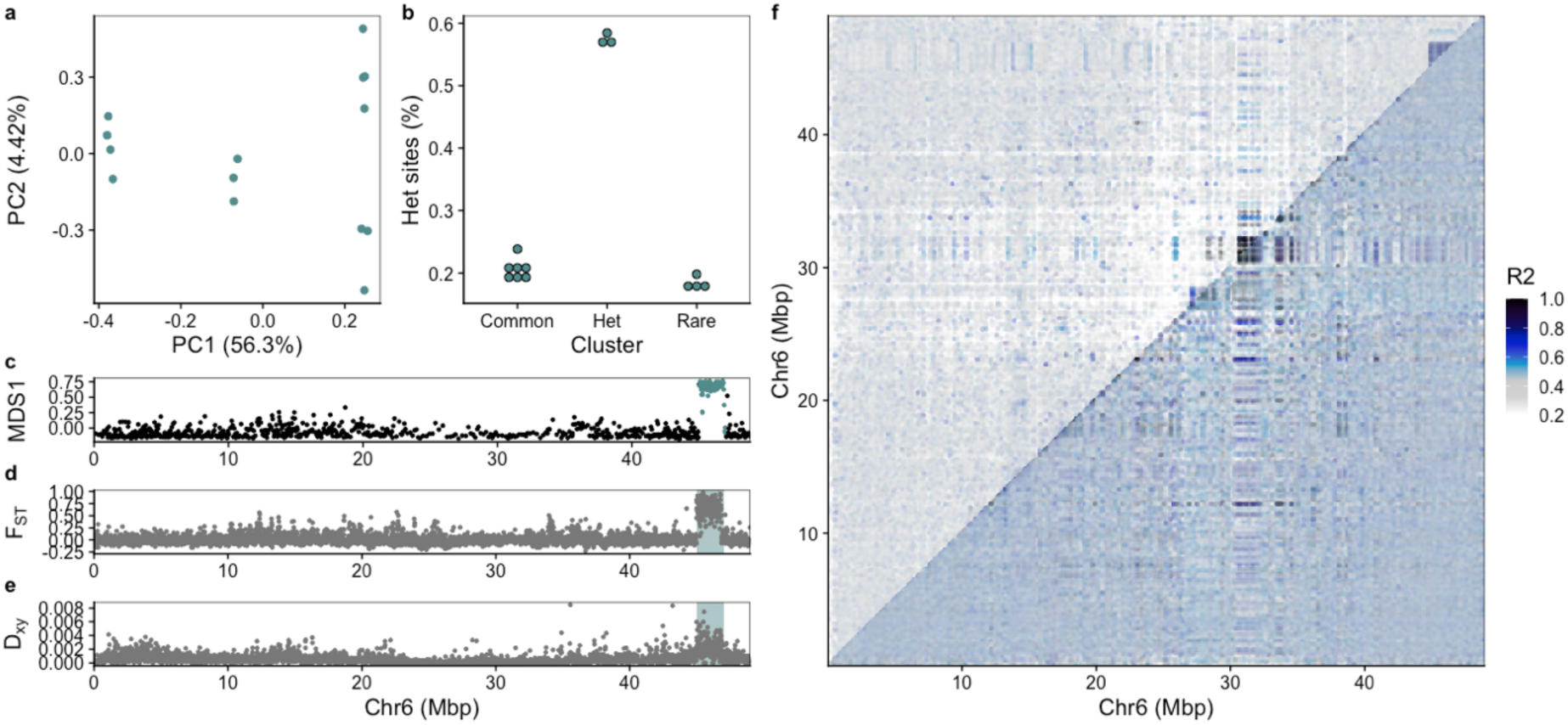
Example of how a chromosomal inversion was detected using chromosome 6 of Arctic cod as reference. a) PCA for the inversion region identified using lostruct. b) Manually assigned cluster groups and heterozygous sites given in bins for the clusters. c) MDS analysis produced by lostruct where the inversion region is highlighted. d) F_ST_ and e) D_XY_ calculated with pixy showing elevated values within the highlighted inversion region. f) pairwise linkage disequilibrium plot calculated using pixy where the top triangle includes all samples, and the lower triangle includes only the individuals within the common type. The upper right corner of the top triangle shows elevated R2 values; however, the bottom triangle, containing only individuals with homokaryotypes of the common type, does not display elevated R2 values. This pattern is in line with what is expected for a chromosomal inversion.

**Table 1:**
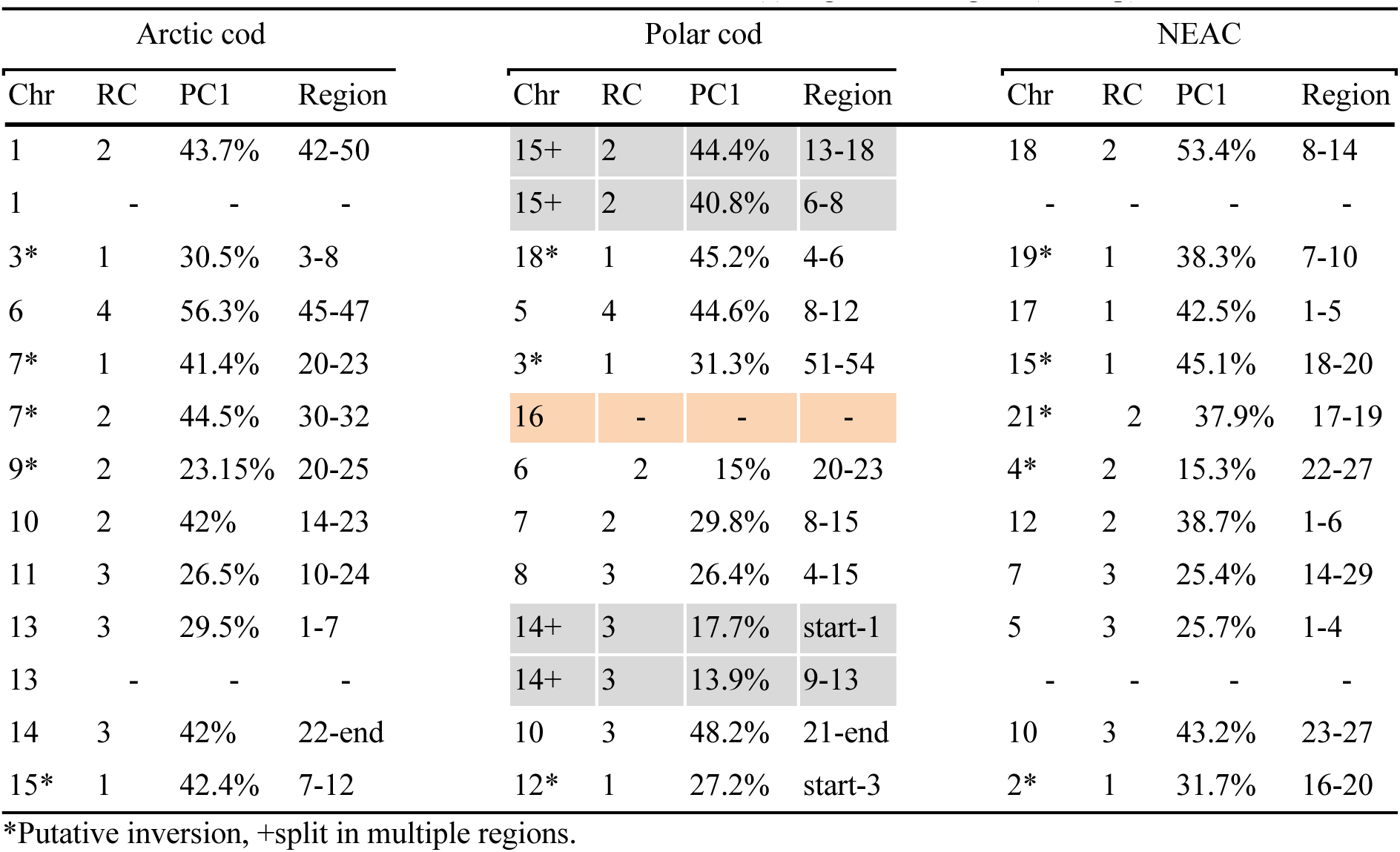
Inferred chromosomal inversions for Arctic cod using the three reference genomes. First column gives chromosome (Chr) in Arctic cod and the homologous chromosome is given for the other two species. The count of individuals is given in the rare group (RC), explained variation for the first principal component (PC1), and the region used to run PCA calculations. The grey coloring indicates the inversion split in two, while orange coloring denotes inversion not detected. Location on the chromosome(s) is given as Region (in Mbp).

| Arctic cod |  |  |  | Polar cod |  |  |  | NEAC |  |  |  |
| --- | --- | --- | --- | --- | --- | --- | --- | --- | --- | --- | --- |
| Chr | RC | PC1 | Region | Chr | RC | PC1 | Region | Chr | RC | PC1 | Region |
| 1 | 2 | 43.7% | 42-50 | 15+ | 2 | 44.4% | 13-18 | 18 | 2 | 53.4% | 8-14 |
| 1 | - | - | - | 15+ | 2 | 40.8% | 6-8 | - | - | - | - |
| 3* | 1 | 30.5% | 3-8 | 18* | 1 | 45.2% | 4-6 | 19* | 1 | 38.3% | 7-10 |
| 6 | 4 | 56.3% | 45-47 | 5 | 4 | 44.6% | 8-12 | 17 | 1 | 42.5% | 1-5 |
| 7* | 1 | 41.4% | 20-23 | 3* | 1 | 31.3% | 51-54 | 15* | 1 | 45.1% | 18-20 |
| 7* | 2 | 44.5% | 30-32 | 16 | - | - | - | 21* | 2 | 37.9% | 17-19 |
| 9* | 2 | 23.15% | 20-25 | 6 | 2 | 15% | 20-23 | 4* | 2 | 15.3% | 22-27 |
| 10 | 2 | 42% | 14-23 | 7 | 2 | 29.8% | 8-15 | 12 | 2 | 38.7% | 1-6 |
| 11 | 3 | 26.5% | 10-24 | 8 | 3 | 26.4% | 4-15 | 7 | 3 | 25.4% | 14-29 |
| 13 | 3 | 29.5% | 1-7 | 14+ | 3 | 17.7% | start-1 | 5 | 3 | 25.7% | 1-4 |
| 13 | - | - | - | 14+ | 3 | 13.9% | 9-13 | - | - | - | - |
| 14 | 3 | 42% | 22-end | 10 | 3 | 48.2% | 21-end | 10 | 3 | 43.2% | 23-27 |
| 15* | 1 | 42.4% | 7-12 | 12* | 1 | 27.2% | start-3 | 2* | 1 | 31.7% | 16-20 |

\*Putative inversion, +split in multiple regions.

The larger number of inversions detected in Arctic cod is comparable with the higher number of inversions detected in polar cod, where in total 20 inversions are detected^28^. Both species resides in freezing water temperatures, and thus it is speculated that this high number is linked to cold water adaptions^27^.

### Reference bias in inversion detection coupled to interspecific chromosomal reshufflings and translocations

By taking full advantage of the *intraspecific* datasets, we uncovered that the precision, in terms of size and location, of the inversion scoring became problematic when using a different reference genome than the focal species (Table 1). Employing NEAC as the reference genome, all six validated inversions were confirmed as well as the putative inversions (Figure S18-S28). Even though all the inversions were detected, the majority of the inversions identified were not found to be of similar size and nor with the same chromosomal positioning as the corresponding inversions detected using Arctic cod as the reference. This differentiation is mainly due to the larger species-specific genomic rearrangements and translocations that have occurred within this lineage (Figure 6, 7 and Hoff et al.^27^). However, for some of the inversions a partly overlapping positioning was detected when inspecting homologous chromosomes, i.e., the inversions on chromosome 4, 7, 10, and 19 in NEAC vs. chromosome 9, 11, 14 and 3 in Arctic cod (Table 1). Moreover, two of the inversions detected (on chromosome 7 and 17) in NEAC were found to be larger than the corresponding inversions detected in Arctic cod (on chromosome 11 and 6). Since it has been shown for these two regions that Atlantic cod has overlapping inversions with Arctic cod^27^, this could imply that the signal of the Atlantic cod inversion could infer with the detection of the true Arctic cod inversion.

**Figure 6:**
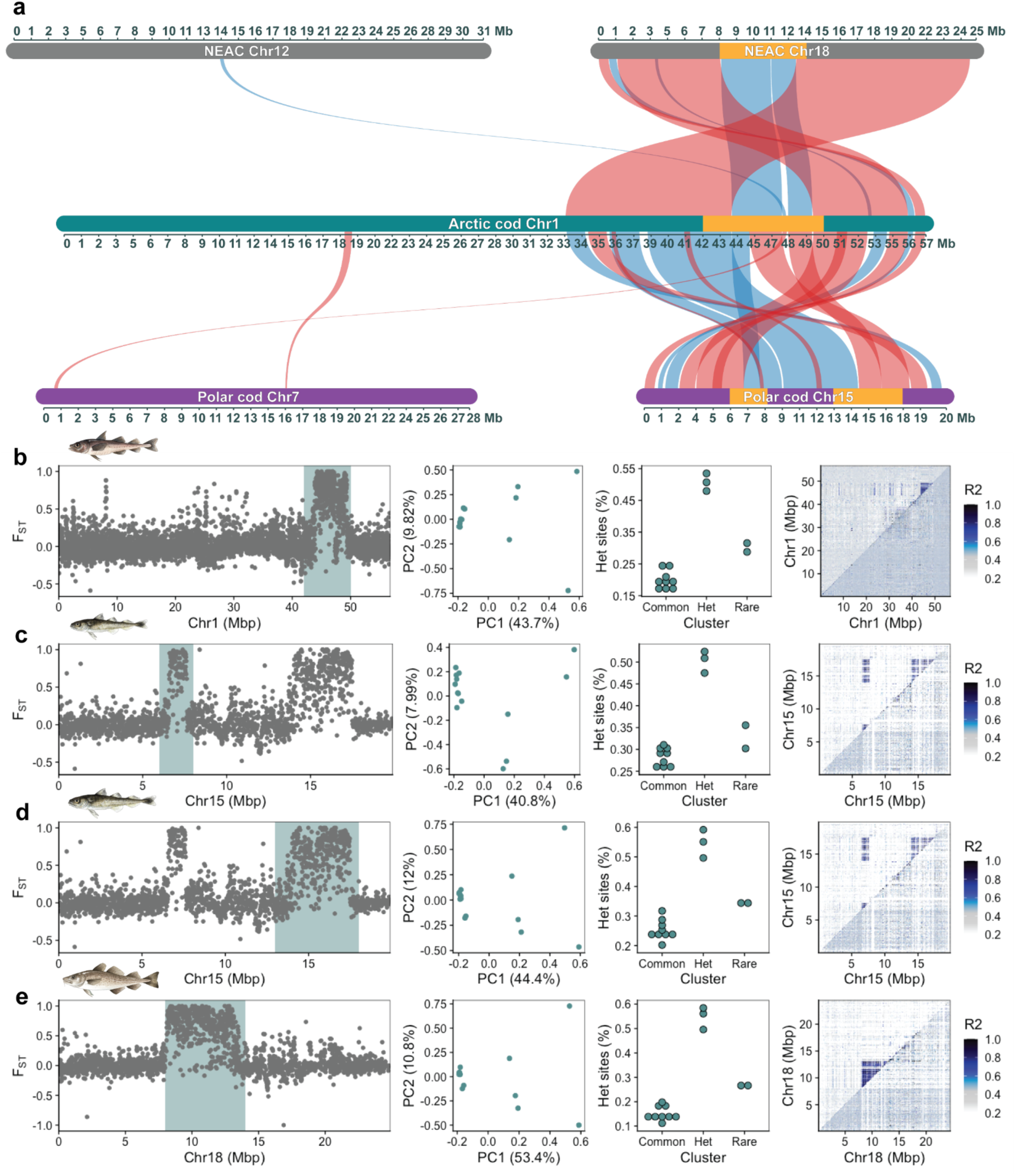
Example of inversion detection bias for the inversion on chromosome 1 in Arctic cod using Arctic cod, polar cod, and NEAC as reference genomes independently. a) Synvisio plots illustrating the structural rearrangements occurring between the three species’ reference genomes for the second half of the Arctic cod chromosome 1; blue indicates the same orientation, while red indicates the reverse orientation, and orange indicates regions defined for the inversion detection protocol. b) Inversion detection when using Arctic cod as a reference. c) and d) Inversion detection when using polar cod as a reference, where the inversion is split into two parts that are linked together. e) When using NEAC as a reference, the inversion is successfully captured. However, a smaller part is missing, as it has translocated to chromosome 12 in NEAC. Each panel is described in further detail in Figure 5.

**Figure 7:**
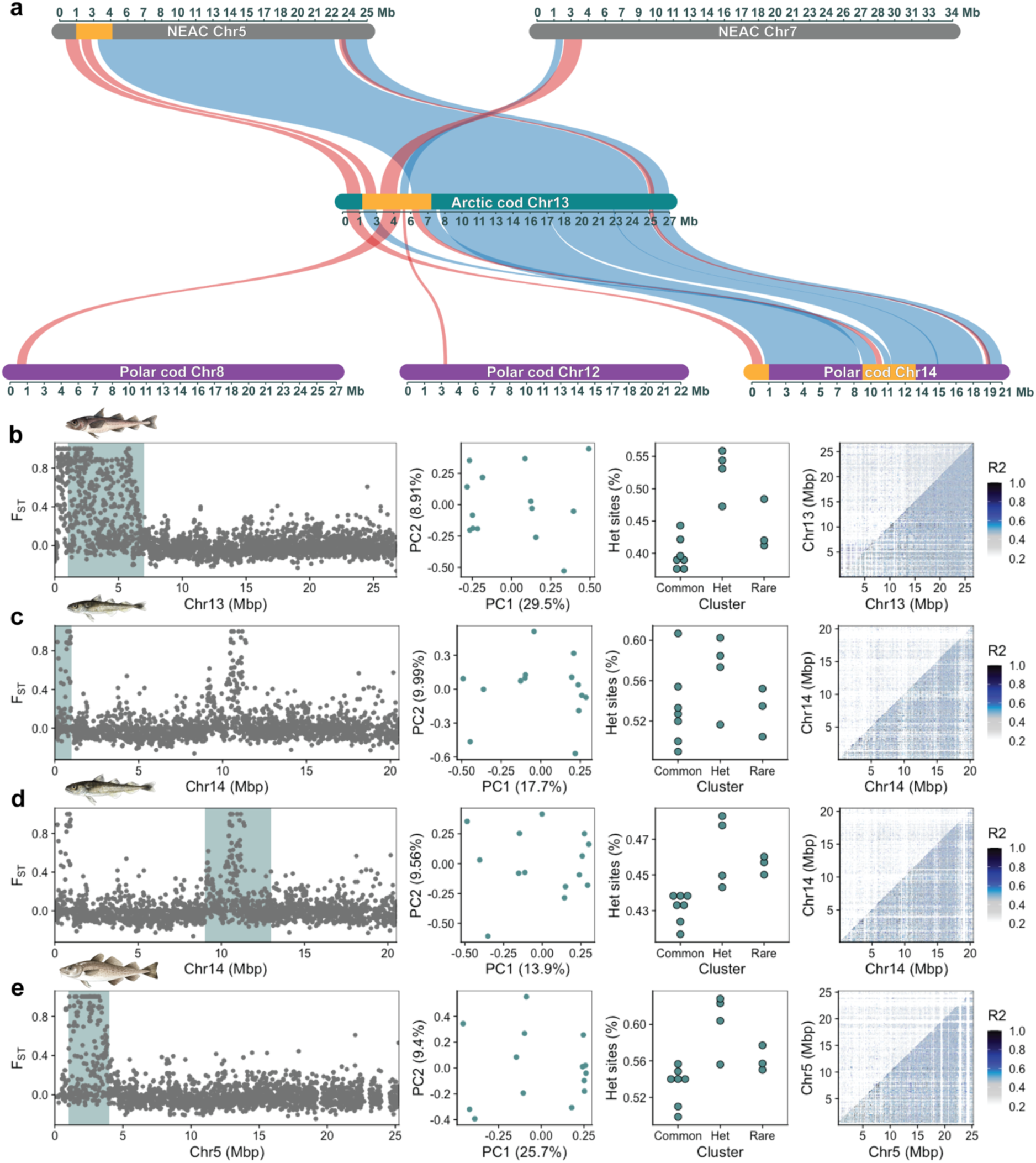
Example of inversion detection bias for the inversion on chromosome 13 in Arctic cod using Arctic cod, polar cod, and NEAC as reference genomes independently. a) Synvisio plots illustrating the structural rearrangements between the three species’ reference genomes for Arctic cod chromosome 13, annotated with the same colors as those used in Figure 5. Here, multiple structural rearrangements between the species obscure the inversion signal for chromosome 13. b) Inversion detection using Arctic cod as a reference. c) and d) Inversion detection using polar cod as a reference, where the inversion appears as two distinct parts. Moreover, the heterozygosity signal is weaker in c), and none of the LD plots capture the inversion when using polar cod as reference. e) The inversion exhibits the expected heterozygosity distribution when using NEAC as a reference, but the LD signal is weak. Each panel is described in further detail in Figure 5.

When applying the more closely related polar cod as the reference, all inversions were detected except one of the putative inversions (Table 1; Figure S29-S41), i.e., the second inversion on chromosome 7 in Arctic cod (corresponding to chromosome 16 in polar cod; Figure S29). Also here, the majority of the inversions identified were not found to be of similar size and nor with the same chromosomal positioning as the corresponding inversions detected using Arctic cod (nor the NEAC) as reference. Moreover, for the inversion detected on chromosome 1 in Arctic cod, the inversion appeared as two separate but linked inversions, a result of chromosomal rearrangements between polar cod and Arctic cod (Figure 6c and d; Figures S30 and S31). Similarly, inaccurate identification due to intraspecific chromosomal translocations (between all three species) is seen for the region enharbouring the inversion on chromosome 13 in Arctic cod (Figure 7). When employing the polar cod genome as the reference, we find that the inversion is split into two separate inversions, with no clear LD signals as well as a less clear heterozygosity distribution (Figure 7c and d; Figure S38 and 39). For the same region using NEAC as reference (Figure 6e; Figure S26), we capture the expected heterozygosity distribution, however only weak LD signals were detected. Adding to the complexity of inversion detection when utilizing a more distantly related refence, the putative inversion on chromosome 3 in Arctic cod showed a much clearer LD signal when either NEAC (Figure S19) or polar cod (Figure S32) was employed.

## Concluding remarks

Our findings combined, strongly indicate that caution must be exercised when using a heterospecific reference genome for variant calling as well as inversion scoring. The quality of the reference used as well as degree of genomic divergence between the focal species and the reference seemingly impact the variants called due to i) a lower degree of mappability and thus, losing informative genetic variation, ii) potential misalignments which could lead to f. ex. a bias towards higher degree of heterozygosity and more noisy datasets, where in-depth analyses on e.g., demography history and detection of signals of selection are highly likely to be erroneous/inflated by this type of reference bias^10–14,61^. Specifically, the general population genetic statistics in terms of heterozygosity, ROH, and genetic diversity, are all metrics that are often used within conservation genomics as measurements for the health situation of a species and/or populations, by estimating the standing genetic variation and thus, their adaptive capacity^4,10,12,62^. Our study shows that some of these metrics are seemingly more sensitive, such as D_XY_ and π, while the F_ST_ estimates are more robust, at least for detecting differentiation between species. However, it could be that F_ST_ estimates may be impacted if looking into differentiation within a species, i.e., between populations and/or ecotypes.

Most importantly, we discovered that the use of reference impacted the detection and characterization of chromosomal inversions. Important information on size, position, and linkage between regions can easily be lost due to species-specific genomic rearrangements and smaller translocations^27^. For instance, when using the more closely related species—the polar cod—as the reference for the detection of inversions in Arctic cod resulted in the detection of several inversions that were defined as two inversions instead of one continuous larger inversion as well one inversion that was not discovered at all. This mismatch in detection of inversions is highly linked to the larger genomic reshufflings that have occurred after Arctic cod and polar cod branched off from their common codfish ancestor ∼4 million years ago^24^. Moreover, most of the inversions detected were smaller than when using the focal species as the reference, i.e., meaning that the breakpoint regions are not fully characterized when using a non-conspecific reference. Additionally, we also uncovered that the precision of detection was impacted if the reference has inversions in the same regions as the focal species. When applying Atlantic cod as the reference, two of the inversions were found to be longer than expected, which could be explained by the fact that Atlantic cod in these regions harbor species-specific but overlapping polymorphic inversions with Arctic cod^21,22,27^. We speculate that highly variable breakpoint regions^63,64^ could lead to higher degree of misalignments in these regions. Taken together, inference of detection of chromosomal inversions when using a non-conspecific reference, should be handled with care. Especially, since the breakpoint regions— where important genes under selection tend to be positioned^27,65–68^—seems to be lost in the scoring of the inversions, as well as the number and interlinking of inversions may be incomplete.

## Author contributions

S.J., S.N.K.H., M.F.M. conceptualized the study. S.N.K.H., M.F.M. did DNA extractions. M.F.M. handled, processed, and analyzed the data. M.R. provided scripts. S.J., S.N.K.H. sampled polar cod and the single specimen of Arctic cod from the Barents Sea. I.B. and K.P. provided Arctic cod specimens from Canada and Greenland, respectively. Funding acquisition by S.J., K.P. and K.S.J.. S.J., S.N.K.H., and M.F.M. did the interpretation and discussion of results. Visualization and design of figures by S.N.K.H., M.F.M., and S.J. J.C. provided early feedback and comments to the manuscript. M.F.M. and S.J. wrote the original manuscript, S.N.K.H. contributed with relevant sections and feedback. All co-authors read, provided feedback, and improved the manuscript.

## Data availability

All unpublished raw sequences from the Arctic cod dataset will be deposited in the European Nucleotide Archive (ENA) at EMBL-EBI upon publication.

## Supporting information

supplementary-materials-ref-bias-arctic-cod

## Acknowledgements

Library preparations and sequencing were performed by the Norwegian Sequencing Centre, Oslo. The computations were performed on resources provided by Sigma2; the National Infrastructure for High Performance Computing and Data Storage in Norway. We thank Alexandra Viertler for the codfish illustrations. We thank the crews of RV Kronprins Haakon (i.e., the Nansen Legacy project) and RV Helmer Hanssen (i.e., the TUNU-cruises) for facilitating the trawling and sample acquisition in the rough Barents Sea and in Northeast Greenland. M.F.M. would like to thank the Nansen Legacy for the early career opportunities provided to him.

## Funding

This work was funded by the Research Council of Norway through the following projects: **‘Nansen Legacy’** (RCN no. 276730) and **‘Comparacod’** (RCN no. 222378).

## Notes

### Competing Interest Statement

The authors have declared no competing interest.

