## supplementary-materials-ref-bias-arctic-cod for "Reference genome bias in light of species-specific chromosomal reorganization and translocations"

#### Table of Contents

**Supplementary Note 1: Sample statistics vs PCA plots of Arctic cod using three references (S1-S3)** 2
**Supplementary Note 2: Mitochondrial demographic inference (S4)** 4
**Workflow for detection of chromosomal inversions (S5)** 7
**Inversion detection plots using Arctic cod as reference genome (S6-S17)** 8
**Inversion detection plots using NEAC as reference genome (S18-S28)** 14
**Inversion detection plots using polar cod as reference genome (S29-S41)** 21
**Supplementary Tables (T1-T4)** 29
**Supplementary Sequencing Report** 30
**Supplementary References** 32

#### Supplementary Note 1: Sample statistics vs PCA plots of Arctic cod using three references (S1-S3)

We used PLINK v1.9[1] to perform a Principal Component Analysis (PCA) on the Arctic cod samples for all three *intraspecific* VCFs. To evaluate potential data biases, we also calculated mean depth, missing sites, and heterozygosity among these samples using VCFtools v0.1.16[2]. These statistics were then plotted alongside the first two principal components to assess statistical bias of samples[3] that might affect the distribution of the samples used.

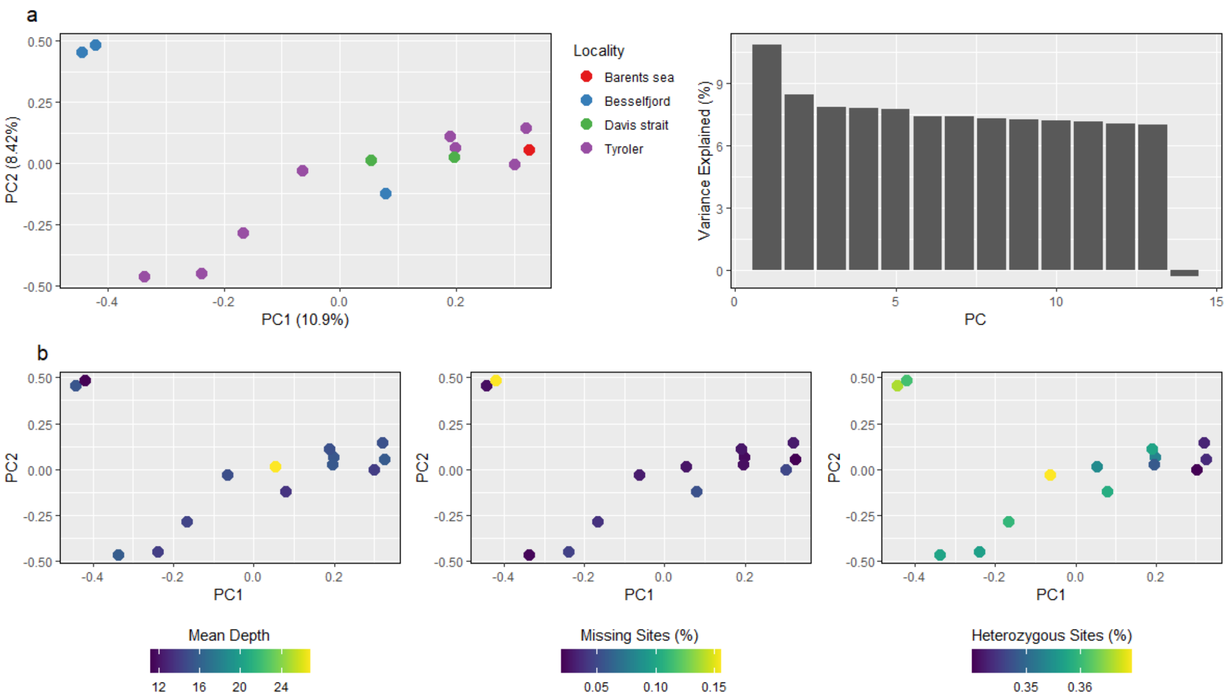


**Supplementary Figure 1:** PCA (PC1 – PC2) versus sample statistics of Arctic cod samples using Arctic cod as reference. a) PCA and corresponding eigenvalues of Arctic cod samples colored by locality. b) PCA against mean depth (left), proportion of missing sites (middle) and proportion of heterozygous sites (right) .

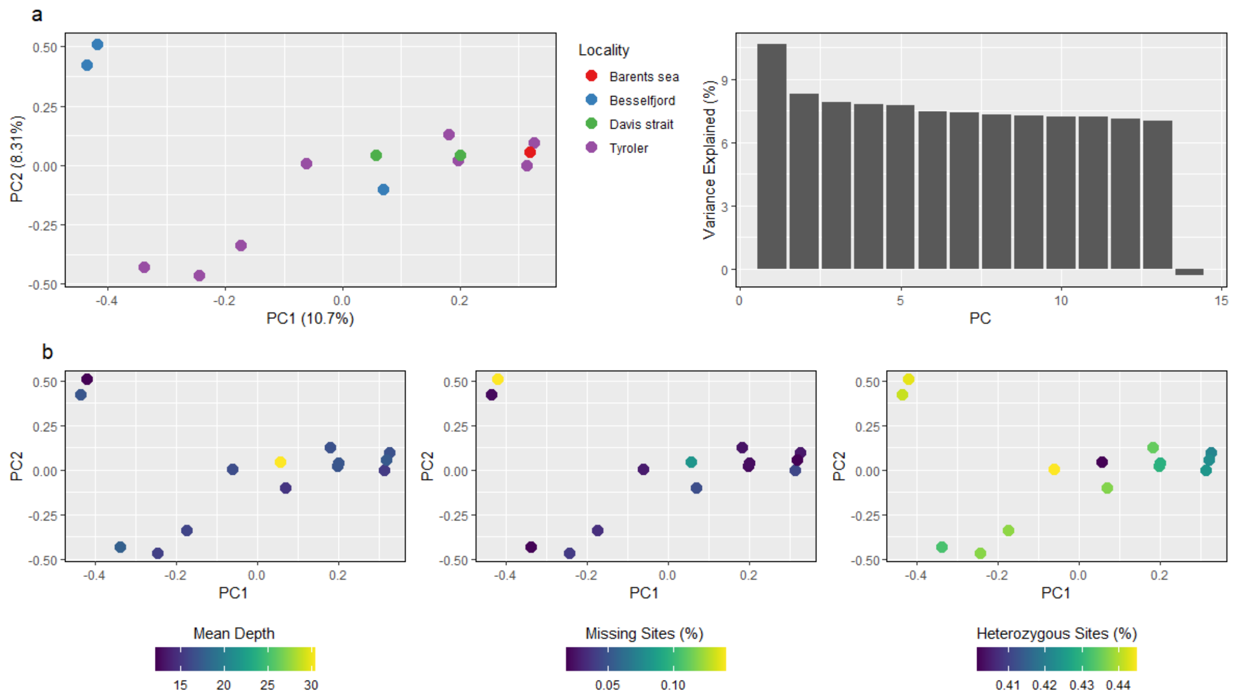


**Supplementary Figure 2:** PCA (PC1 – PC2) versus sample statistics of Arctic cod samples using polar cod as reference. a) PCA and corresponding eigenvalues of Arctic cod samples colored by locality. b) PCA against mean depth (left), proportion of missing sites (middle), and proportion of heterozygous sites (right).

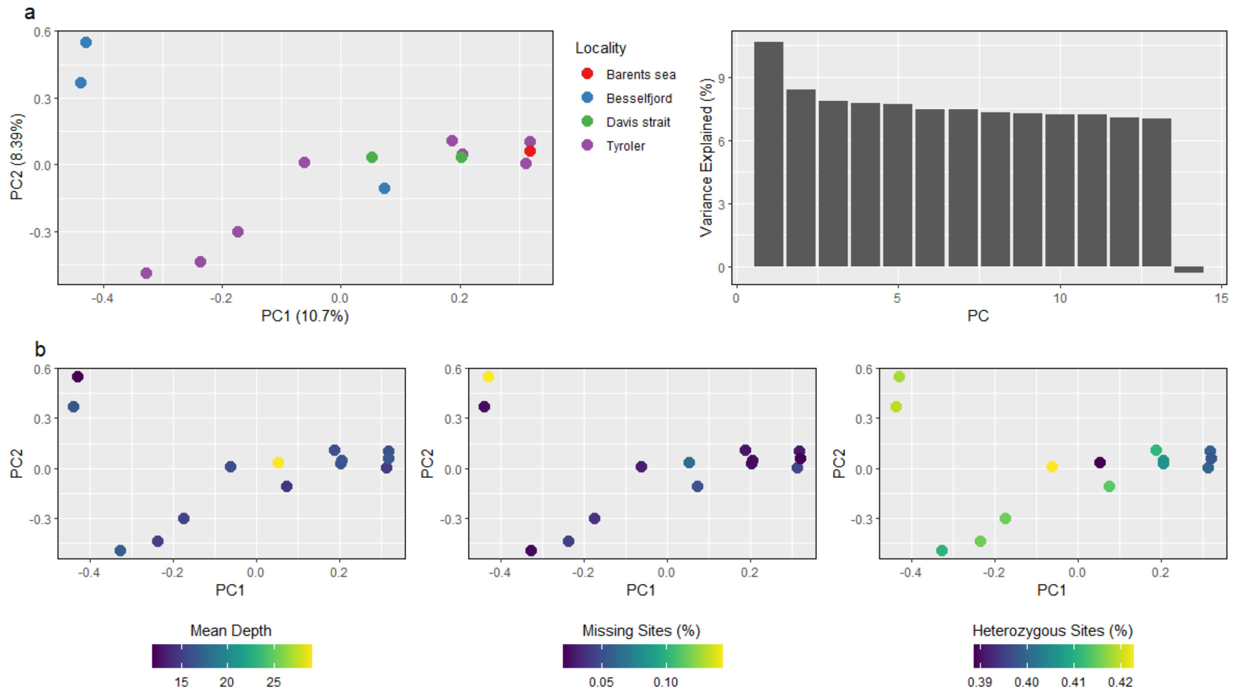


**Supplementary Figure 3:** PCA (PC1 – PC2) versus sample statistics of Arctic cod samples when using NEAC as reference. a) PCA and corresponding eigenvalues of Arctic cod samples colored by locality. b) PCA against mean depth (left), proportion of missing sites (middle) and proportion of heterozygous sites (right).

#### Supplementary Note 2: Mitochondrial demographic inference (S4)

Mitochondrial reads mapped in proper pairs against the Arctic cod[4] mitogenome for each sample were extracted and sorted by read name using SAMtools v1.9[5]. Bam files were converted back to FASTQ format using BEDTools v2.27.1[6] bamtofastq tool. Assembly of mitogenomes was performed using MitoFinder v1.4.1[7] with default settings and the assembler MEGAHIT v1.0[8,9]. MiTFi v0.1[10] was used to annotate the mitochondrial tRNAs. Mitogenome annotation was performed using the available mitogenome reference for Arctic cod (NCBI RefSeq: NC_010122.1) as MitoFinder requires annotated mitogenomes in GenBank reference format. Annotated mitochondrial protein-coding genes (PCGs) from each individually MitoFinder assembled intraspecific level mitogenome of Arctic cod were extracted and aligned using MAFFT v7.453[11]. To allow for all samples to be included, Cytochrome c oxidase subunit II (*COX2*) had to be removed as this gene was missing for one sample. Furthermore, the PCGs were manually corrected for reading frame before they were concatenated with PhyKIT v1.11.7[12] create_concat to produce a supermatrix.

The female effective population size ($N_e$) was estimated in BEAST v2.6.7[13] under the Bayesian skyline model[14]. The substitution model was inferred using bModelTest with the namedExtended list of models. The coalescent Bayesian skyline prior was applied under a strict clock with a rate of $1.14 \times 10^{-8}$ substitution/site/year used, as reported for Atlantic cod[15]. The analysis was performed using a chain length of 800,000,000, and sampling was done every 1,000 iterations with bPopSizes and bGroupSizes set to 5 dimensions. Tracer v1.7.2 was used to check convergence and to reconstruct the Bayesian skyline with default settings. (skyline variant = stepwise (constant), maximum time to root height = lower 95% HPD, root height = Treeheight, number of bins = 100).

Due to the low sample count of Arctic cod, Bayesian skyline analysis was repeated while including the available Arctic cod samples from NCBI. Samples sourced from NCBI included 19 Arctic cod samples from Wilson et al[16]. and one Arctic cod sample from Breines et al[17]. These mitogenomes were in partial condition. Therefore, to maximize sample count and PCG count, the individual with missing *COX2* was removed, resulting in a total of 33 samples and 10 PCGs.

Demographic inference based on the 14 Arctic cod individuals (Figure S4a) indicated an increase in female effective population size ($N_e$) around the last glacial maximum (LGM), as well as a slight diagonal increase around 40 Kya. When Arctic cod samples from NCBI were included (N=33, Figure S4b), the $N_e$ increase was steepest around 40 Kya, with a slight diagonal increase after the LGM.

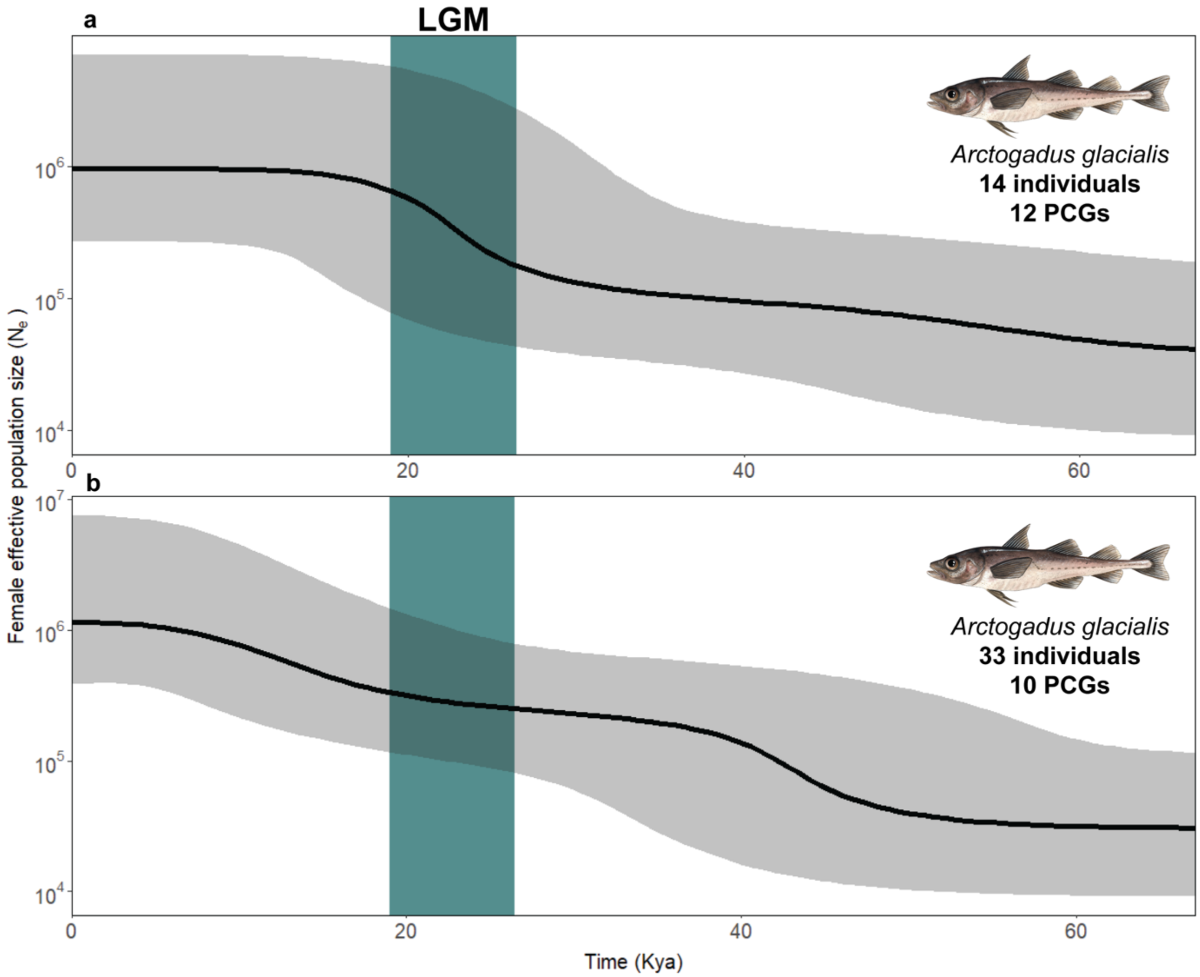


**Supplementary Figure 4:** Inferred female effective population size ($N_e$) for Arctic cod and mitochondrial demographic inference. a) Demographic history of Arctic using the 14 individuals from this study and 12 PCGs (*COX2* removed due to it being incomplete within one individual), b) Arctic cod including 20 samples sourced from NCBI while excluding the individual with incomplete *COX2*. The blue bar indicates the last glacial maximum.

#### Workflow for detection of chromosomal inversions (S5)

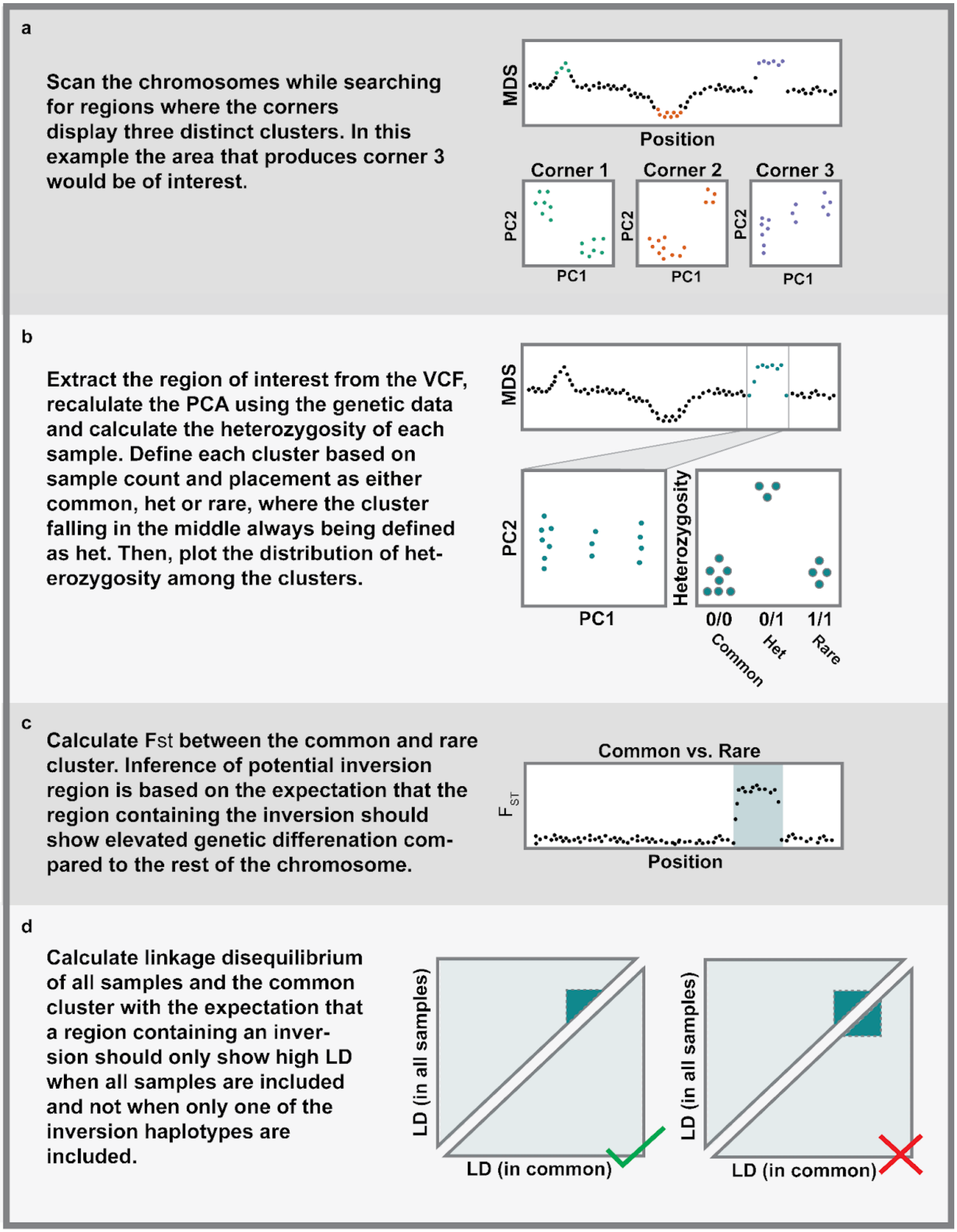


**Supplementary Figure 5:** Illustration of the protocol used to detect chromosomal inversion Arctic cod. The procedure is followed stepwise from a-d.

### Inversion detection plots using Arctic cod as reference genome (S6-S17)

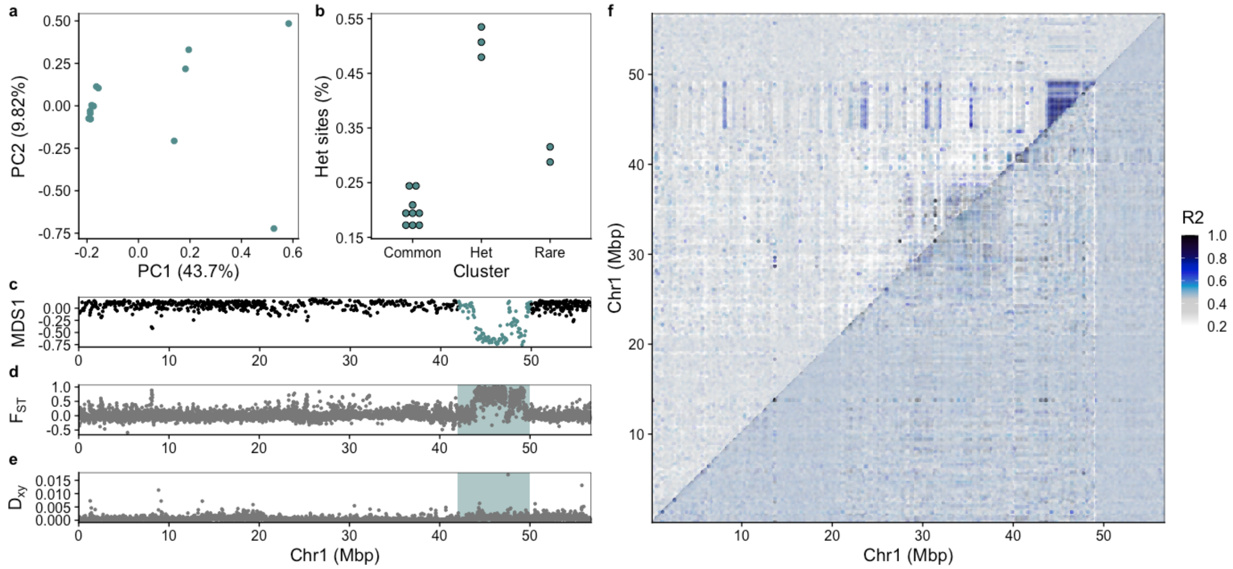


**Supplementary Figure 6:** Inversion detected on chromosome 1 of Arctic cod using Arctic cod as reference genome. a) PCA for the inversion region identified using lostruct. b) Manually assigned cluster groups and heterozygous sites given in bins for the clusters. c) MDS analysis produced by lostruct where the inversion region is highlighted. d) $F_{ST}$ and e) $D_{XY}$ calculated with pixy showing elevated values within the highlighted inversion region. f) pairwise linkage disequilibrium plot calculated using pixy where the top triangle includes all samples, and the lower triangle includes only the individuals within the common type.

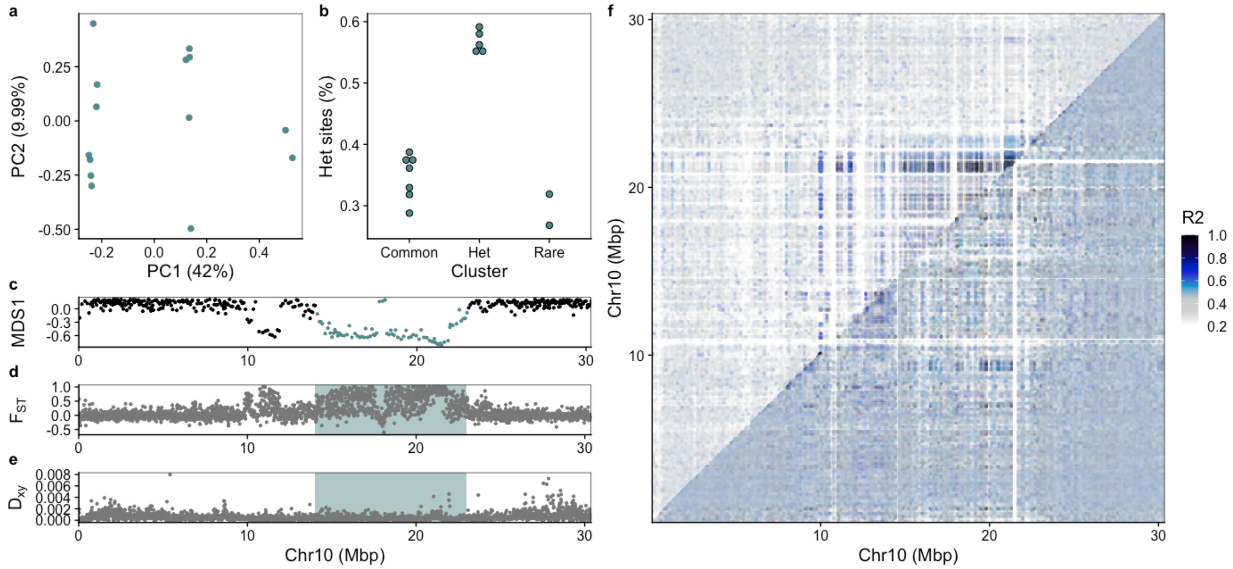


**Supplementary Figure 7:** Inversion detected on chromosome 10 of Arctic cod using Arctic cod as reference genome. a) PCA for the inversion region identified using lostruct. b) Manually assigned cluster groups and heterozygous sites given in bins for the clusters. c) MDS analysis produced by lostruct where the inversion region is highlighted. d) $F_{ST}$ and e) $D_{XY}$ calculated with pixy showing elevated values within the highlighted inversion region. f) pairwise linkage disequilibrium plot calculated using pixy where the top triangle includes all samples, and the lower triangle includes only the individuals within the common type.

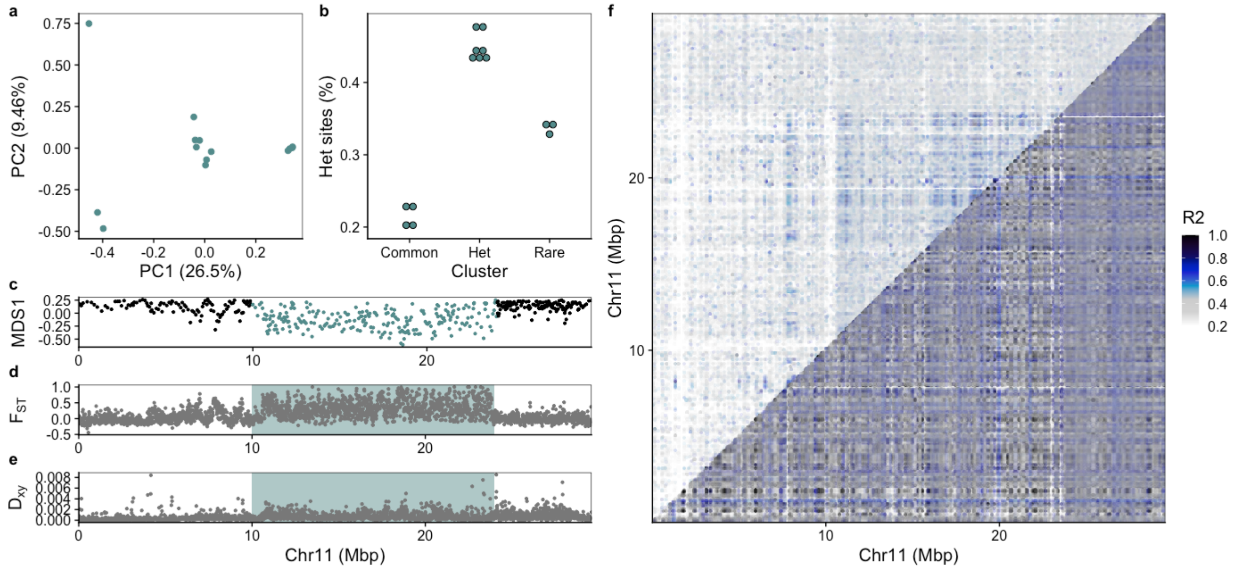


**Supplementary Figure 8:** Inversion detected on chromosome 11 of Arctic cod using Arctic cod as reference genome. a) PCA for the inversion region identified using lostruct. b) Manually assigned cluster groups and heterozygous sites given in bins for the clusters. c) MDS analysis produced by lostruct where the inversion region is highlighted. d) $F_{ST}$ and e) $D_{XY}$ calculated with pixy showing elevated values within the highlighted inversion region. f) pairwise linkage disequilibrium plot calculated using pixy where the top triangle includes all samples, and the lower triangle includes only the individuals within the common type.

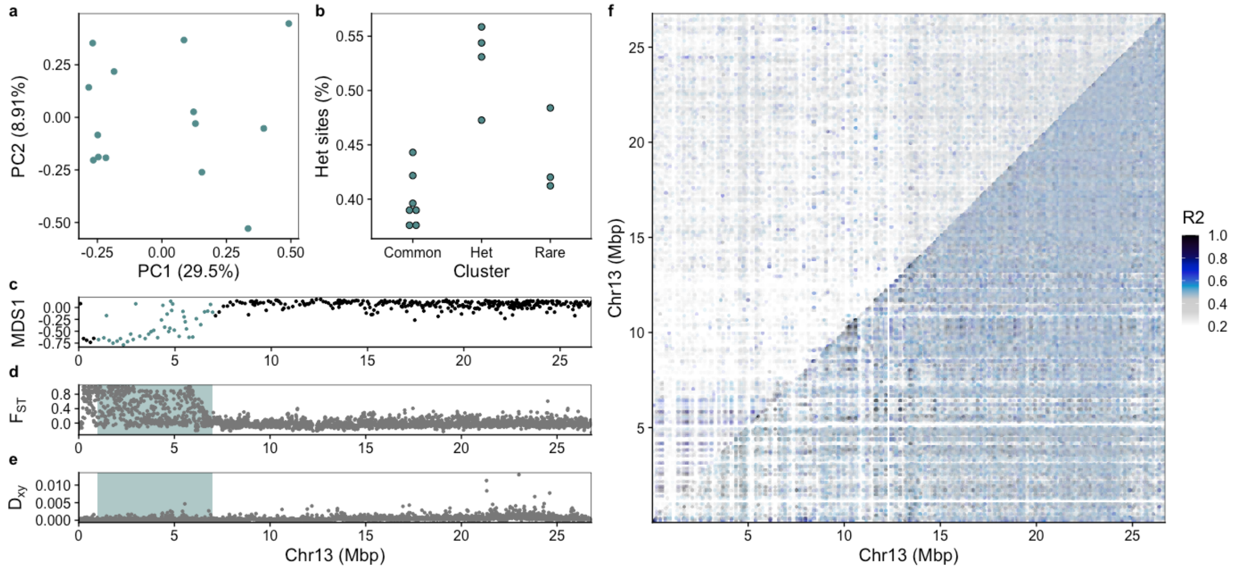


**Supplementary Figure 9:** Inversion detected on chromosome 13 of Arctic cod using Arctic cod as reference genome. a) PCA for the inversion region identified using lostruct. b) Manually assigned cluster groups and heterozygous sites given in bins for the clusters. c) MDS analysis produced by lostruct where the inversion region is highlighted. d) $F_{ST}$ and e) $D_{XY}$ calculated with pixy showing

elevated values within the highlighted inversion region. f) pairwise linkage disequilibrium plot calculated using pixy where the top triangle includes all samples, and the lower triangle includes only the individuals within the common type.

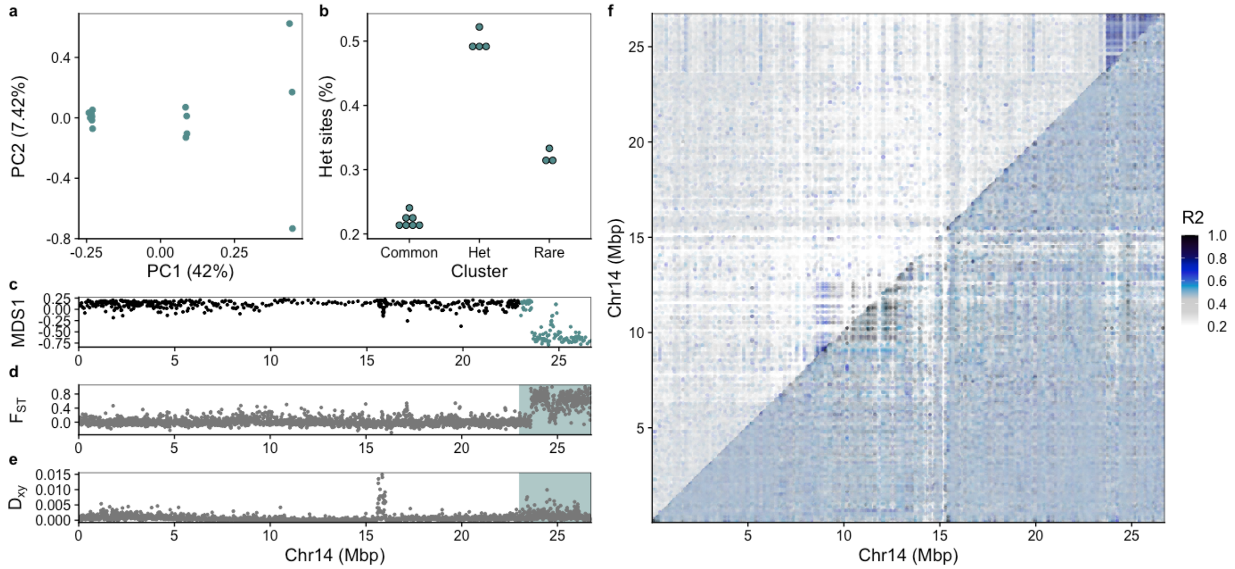


**Supplementary Figure 10:** Inversion detected on chromosome 14 of Arctic cod using Arctic cod as reference genome. See Figure 5 in the main text for descriptions of the different panels. a) PCA for the inversion region identified using lostruct. b) Manually assigned cluster groups and heterozygous sites given in bins for the clusters. c) MDS analysis produced by lostruct where the inversion region is highlighted. d) $F_{ST}$ and e) $D_{XY}$ calculated with pixy showing elevated values within the highlighted inversion region. f) pairwise linkage disequilibrium plot calculated using pixy where the top triangle includes all samples, and the lower triangle includes only the individuals within the common type.

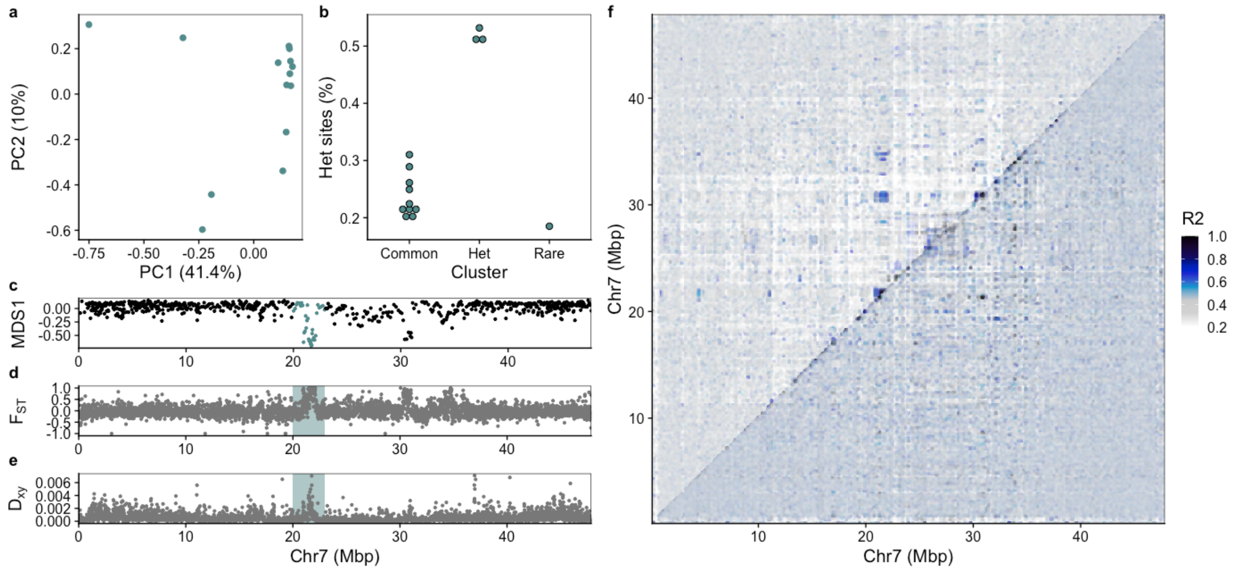


**Supplementary Figure 11:** Putative inversion detected on chromosome 7 (1) of Arctic cod using Arctic cod as reference genome. a) PCA for the inversion region identified using lostruct. b) Manually assigned cluster groups and heterozygous sites given in bins for the clusters. c) MDS analysis produced by lostruct where the inversion region is highlighted. d) $F_{ST}$ and e) $D_{XY}$ calculated with pixy showing elevated values within the highlighted inversion region. f) pairwise linkage disequilibrium plot calculated using pixy where the top triangle includes all samples, and the lower triangle includes only the individuals within the common type.

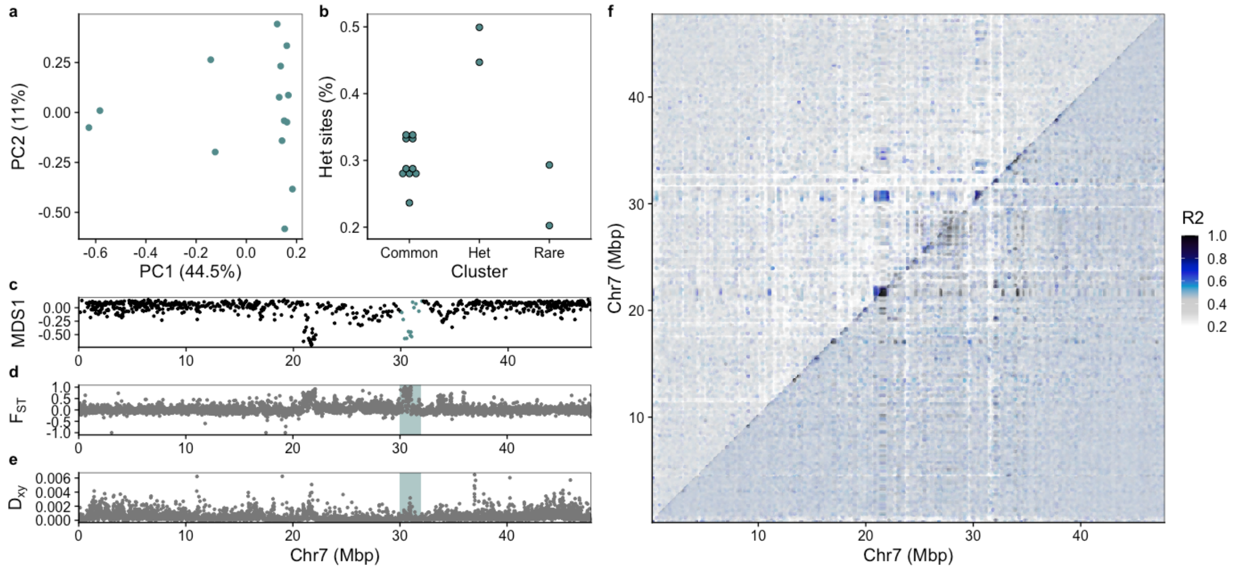


**Supplementary Figure 12:** Putative inversion detected on chromosome 7 (2) of Arctic cod using Arctic cod as reference genome. a) PCA for the inversion region identified using lostruct. b) Manually assigned cluster groups and heterozygous sites given in bins for the clusters. c) MDS analysis produced by lostruct where the inversion region is highlighted. d) $F_{ST}$ and e) $D_{XY}$ calculated with pixy showing elevated values within the highlighted inversion region. f) pairwise linkage disequilibrium plot calculated using pixy where the top triangle includes all samples, and the lower triangle includes only the individuals within the common type.

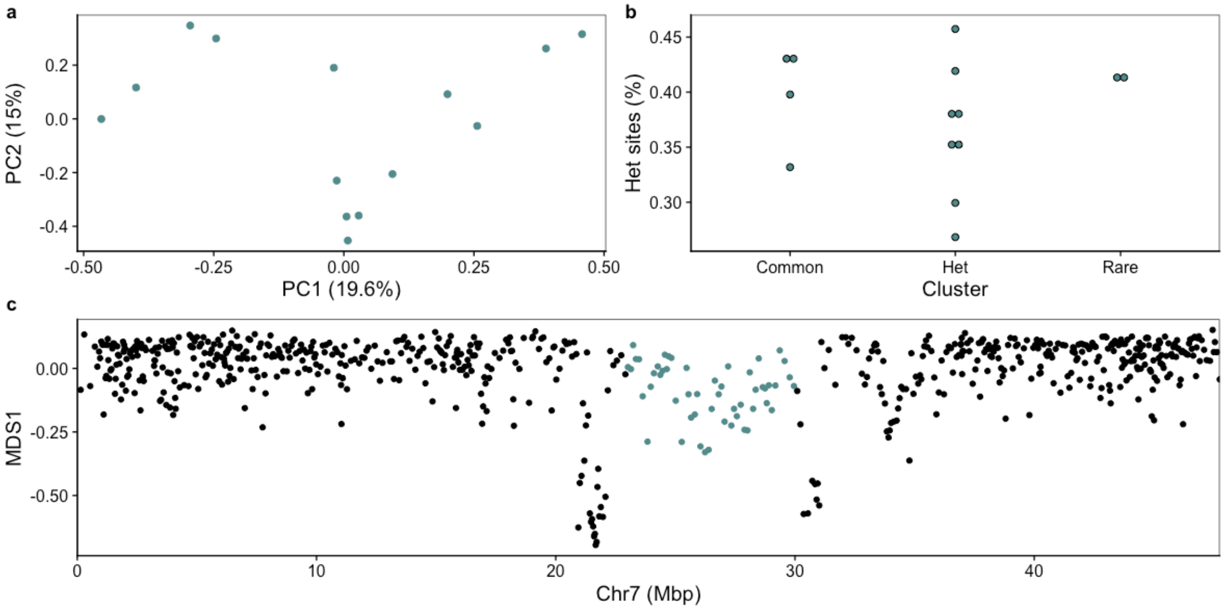


**Supplementary Figure 13:** Testing for inversion signal between the two putative inversions on chromosome 7 of Arctic cod using Arctic cod as reference genome. a) PCA of the region does not show an inversion signal, b) clusters do not follow the typical inversion heterozygosity pattern, and c) MDS1 from lostruct where the region used for calculating the PCA is highlighted.

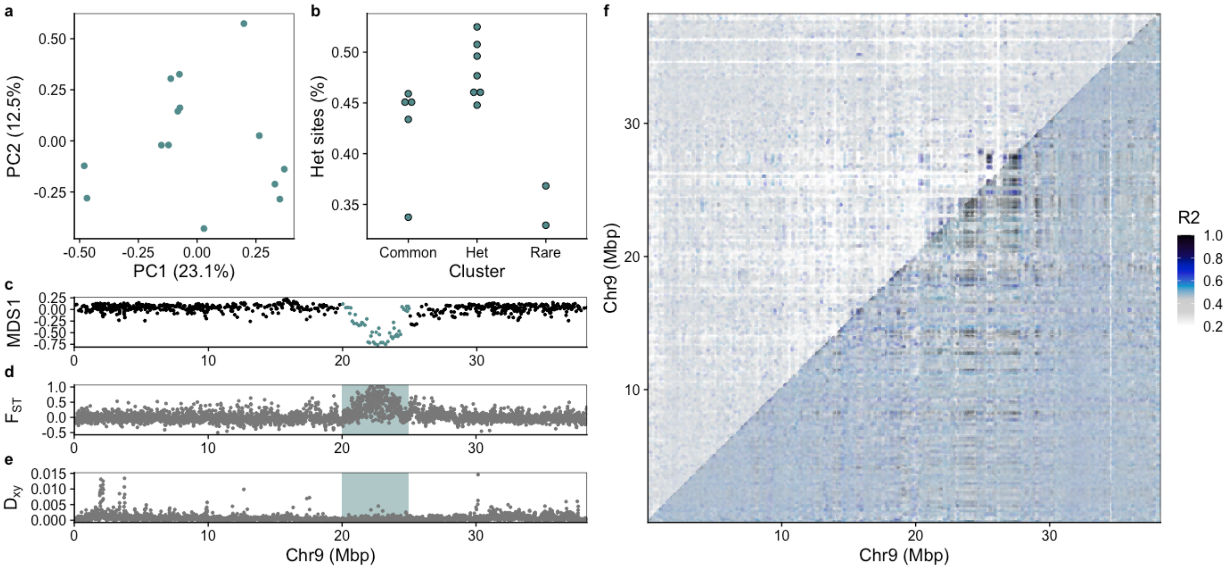


**Supplementary Figure 14:** Putative inversion detected on chromosome 9 of Arctic cod using Arctic cod as reference genome. a) PCA for the inversion region identified using lostruct. b) Manually assigned cluster groups and heterozygous sites given in bins for the clusters. c) MDS analysis produced by lostruct where the inversion region is highlighted. d) $F_{ST}$ and e) $D_{XY}$ calculated with pixy showing elevated values within the highlighted inversion region. f) pairwise linkage disequilibrium plot calculated using pixy where the top triangle includes all samples, and the lower triangle includes only the individuals within the common type.

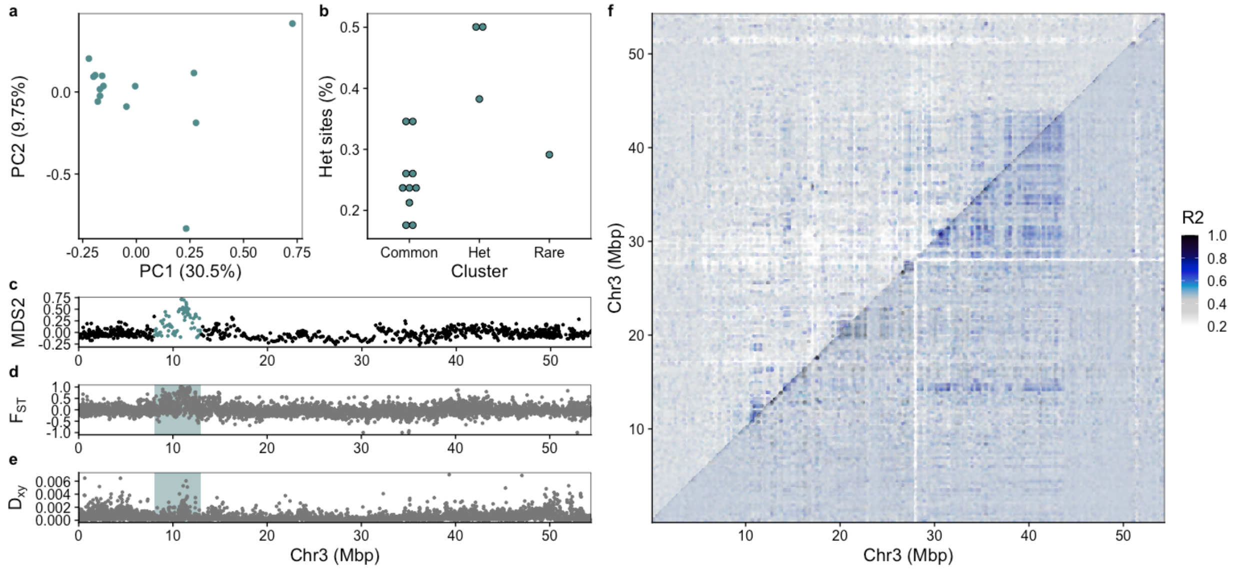


**Supplementary Figure 15:** Putative inversion detected on chromosome 3 of Arctic cod using Arctic cod as reference genome. a) PCA for the inversion region identified using lostruct. b) Manually assigned cluster groups and heterozygous sites given in bins for the clusters. c) MDS analysis produced by lostruct where the inversion region is highlighted. d) $F_{ST}$ and e) $D_{XY}$ calculated with pixy showing elevated values within the highlighted inversion region. f) pairwise linkage disequilibrium plot calculated using pixy where the top triangle includes all samples, and the lower triangle includes only the individuals within the common type.

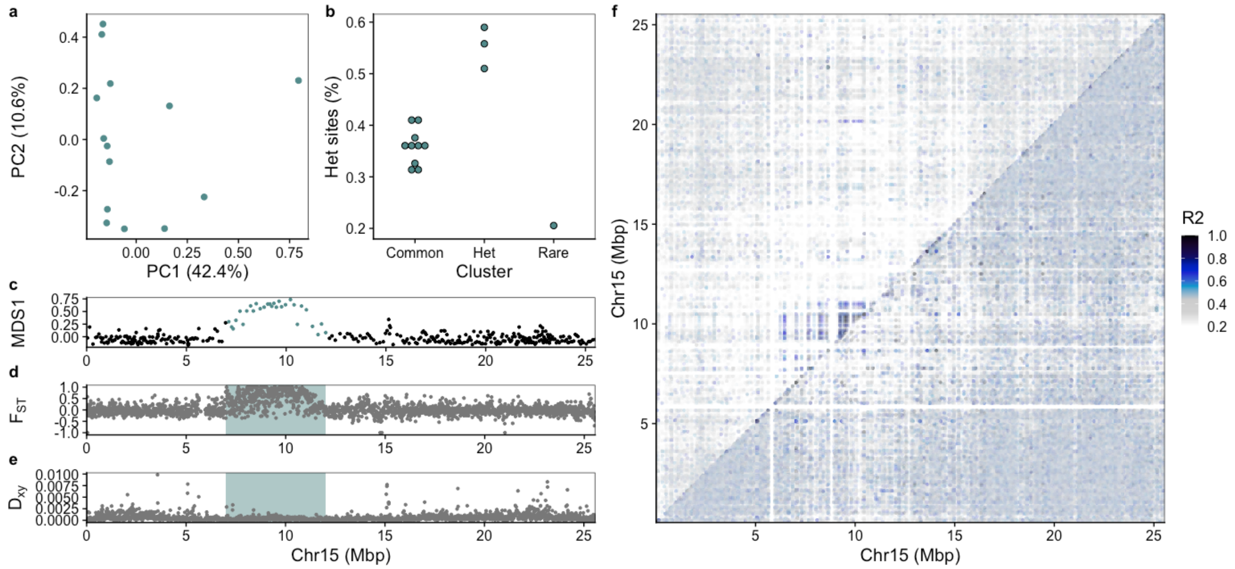


**Supplementary Figure 16:** Putative inversion detected on chromosome 15 of Arctic cod using Arctic cod as reference genome. a) PCA for the inversion region identified using lostruct. b) Manually assigned cluster groups and heterozygous sites given in bins for the clusters. c) MDS analysis produced by lostruct where the inversion region is highlighted. d) $F_{ST}$ and e) $D_{XY}$ calculated with pixy showing elevated values within the highlighted inversion region. f) pairwise linkage disequilibrium plot calculated using pixy where the top triangle includes all samples, and the lower triangle includes only the individuals within the common type.

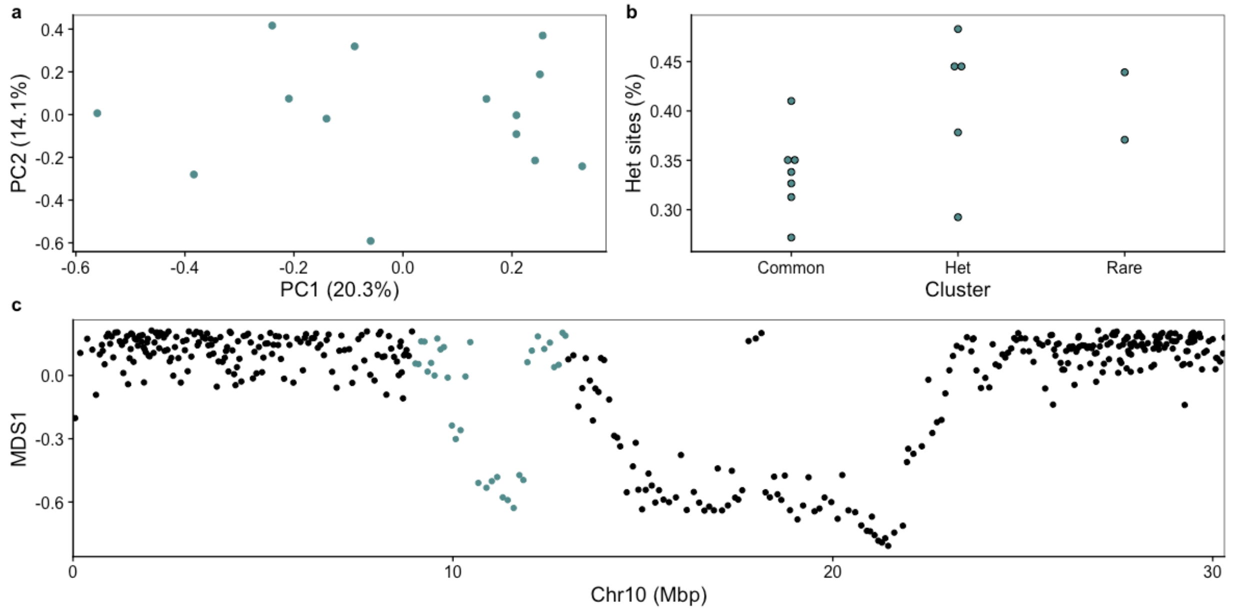


**Supplementary Figure 17:** Testing for inversion signal for a region of differentiation before the inversion of chromosome 10 of Arctic cod using Arctic cod as reference genome. a) PCA of the region does not show a clear inversion signal, b) clusters have a weak heterozygosity pattern, and c) MDS1 from lostruct where the region used for calculating the PCA is highlighted.

### Inversion detection plots using NEAC as reference genome (S18-S28)

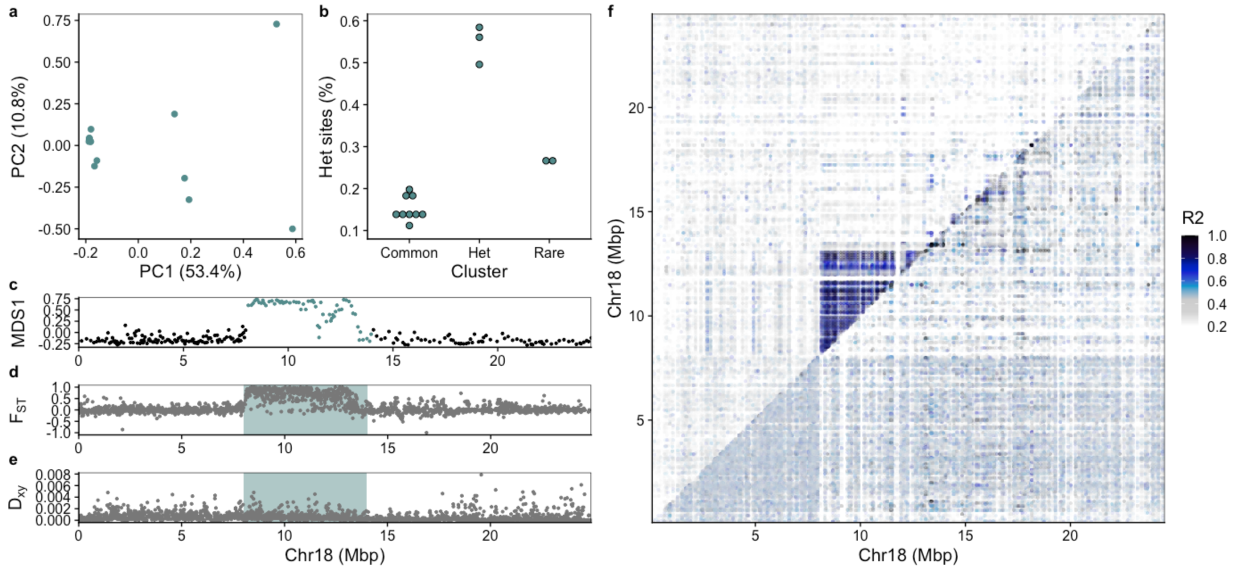


**Supplementary Figure 18:** Inversion detected on chromosome 18 (Arctic cod Chr1) using NEAC as reference genome. a) PCA for the inversion region identified using lostruct. b) Manually assigned cluster groups and heterozygous sites given in bins for the clusters. c) MDS analysis produced by lostruct where the inversion region is highlighted. d) $F_{ST}$ and e) $D_{XY}$ calculated with pixy showing elevated values within the highlighted inversion region. f) pairwise linkage disequilibrium plot calculated using pixy where the top triangle includes all samples, and the lower triangle includes only the individuals within the common type.

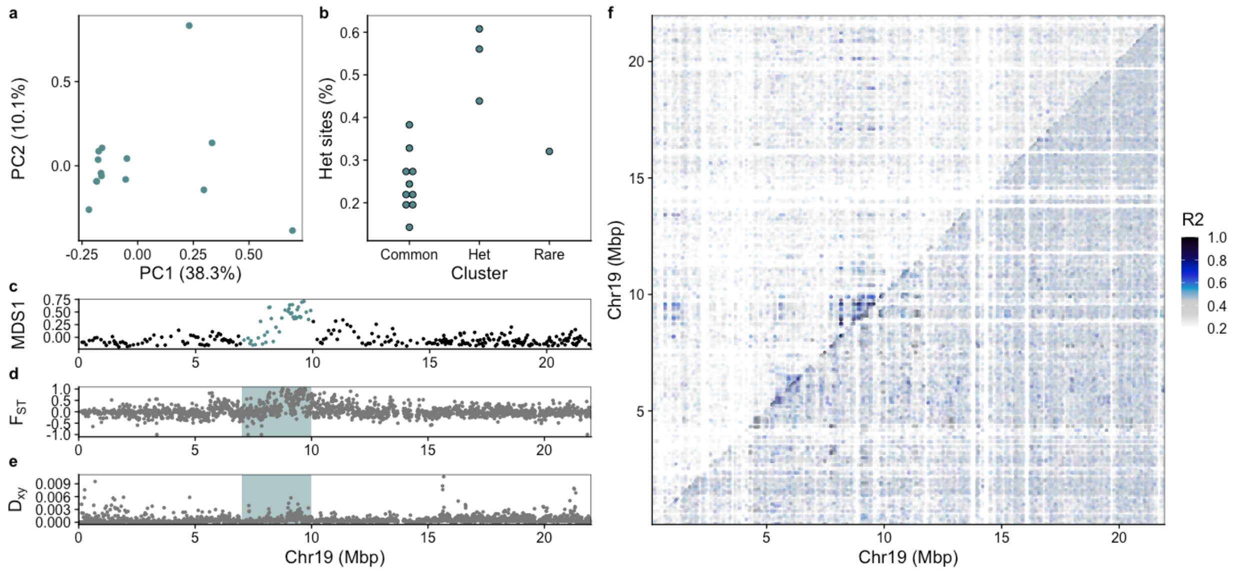


**Supplementary Figure 19:** Putative inversion detected on chromosome 19 (Arctic cod Chr3) using NEAC as reference genome. a) PCA for the inversion region identified using lostruct. b) Manually assigned cluster groups and heterozygous sites given in bins for the clusters. c) MDS analysis produced by lostruct where the inversion region is highlighted. d) $F_{ST}$ and e) $D_{XY}$ calculated with pixy showing elevated values within the highlighted inversion region. f) pairwise linkage disequilibrium plot calculated using pixy where the top triangle includes all samples, and the lower triangle includes only the individuals within the common type.

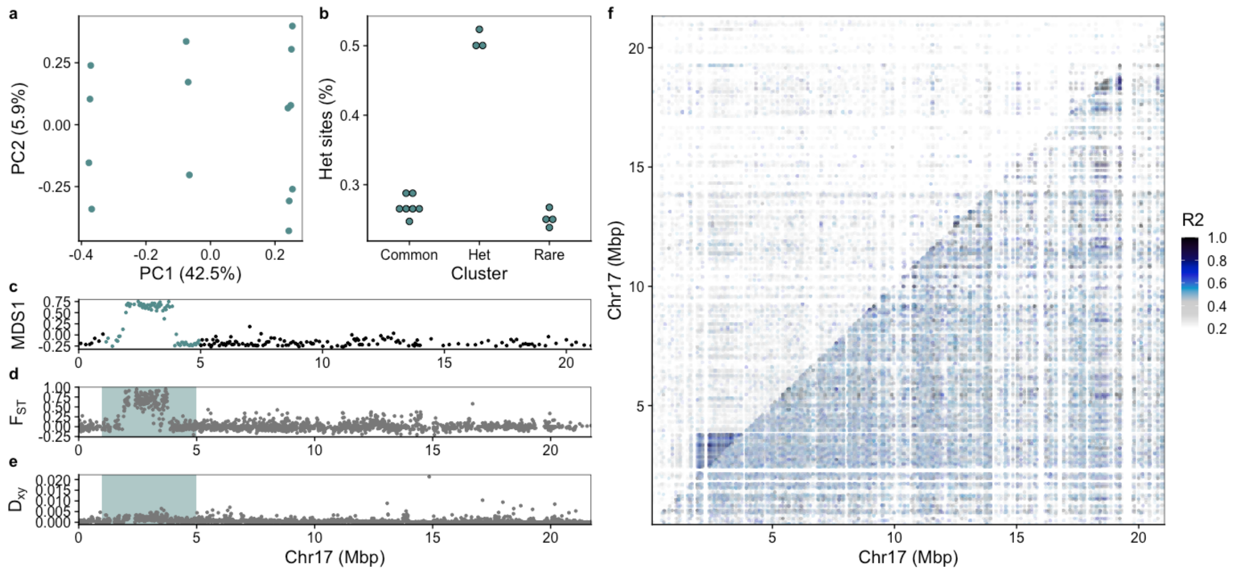


**Supplementary Figure 20:** Inversion detected on chromosome 17 (Arctic cod Chr6) using NEAC as reference genome. a) PCA for the inversion region identified using lostruct. b) Manually assigned cluster groups and heterozygous sites given in bins for the clusters. c) MDS analysis produced by lostruct where the inversion region is highlighted. d) $F_{ST}$ and e) $D_{XY}$ calculated with pixy showing

elevated values within the highlighted inversion region. f) pairwise linkage disequilibrium plot calculated using pixy where the top triangle includes all samples, and the lower triangle includes only the individuals within the common type.

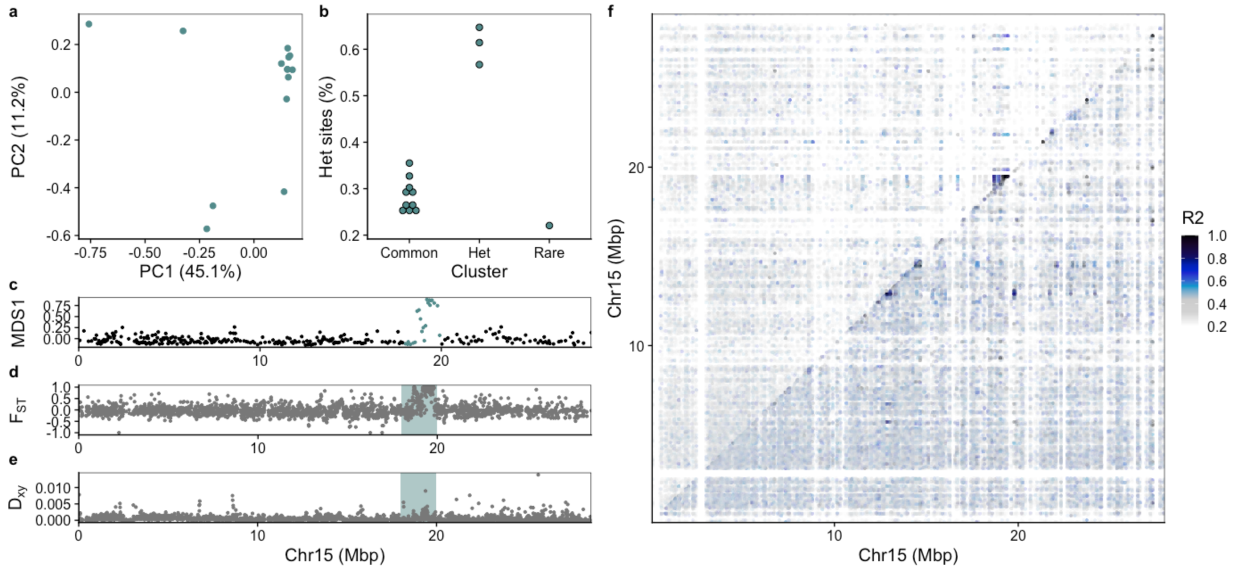


**Supplementary Figure 21:** Putative inversion detected on chromosome 15 (Arctic cod Chr7 (1)) using NEAC as reference genome. a) PCA for the inversion region identified using lostruct. b) Manually assigned cluster groups and heterozygous sites given in bins for the clusters. c) MDS analysis produced by lostruct where the inversion region is highlighted. d) $F_{ST}$ and e) $D_{XY}$ calculated with pixy showing elevated values within the highlighted inversion region. f) pairwise linkage disequilibrium plot calculated using pixy where the top triangle includes all samples, and the lower triangle includes only the individuals within the common type.

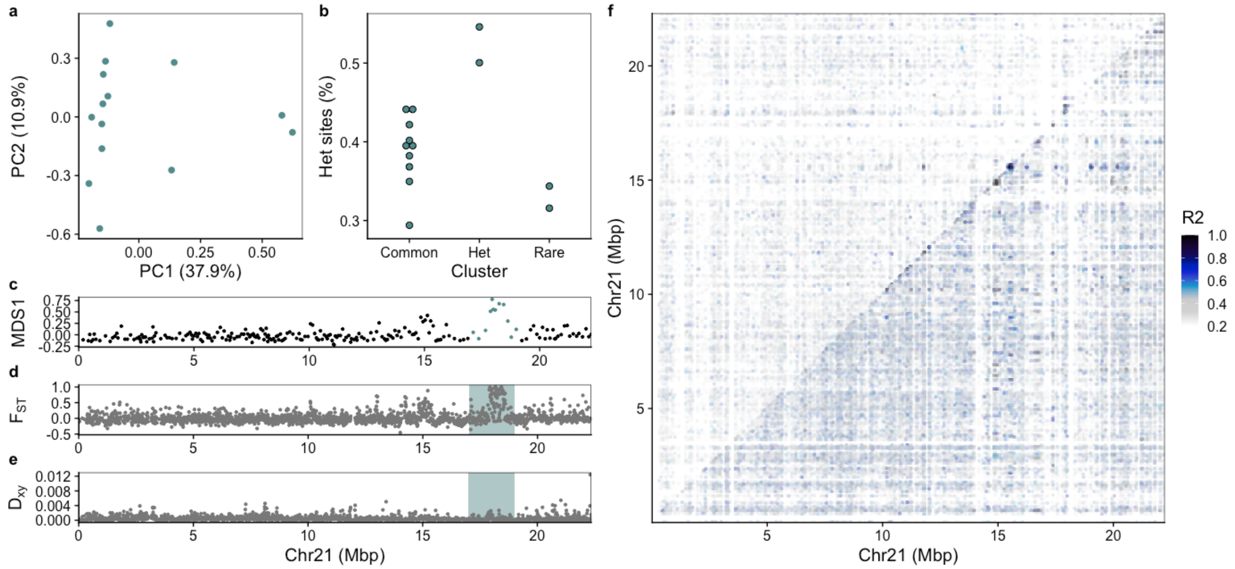


**Supplementary Figure 22:** Putative inversion detected on chromosome 21 (Arctic cod Chr7 (2)) using NEAC as reference genome. a) PCA for the inversion region identified using lostruct. b) Manually assigned cluster groups and heterozygous sites given in bins for the clusters. c) MDS analysis produced by lostruct where the inversion region is highlighted. d) $F_{ST}$ and e) $D_{XY}$ calculated with pixy showing elevated values within the highlighted inversion region. f) pairwise linkage disequilibrium plot calculated using pixy where the top triangle includes all samples, and the lower triangle includes only the individuals within the common type.

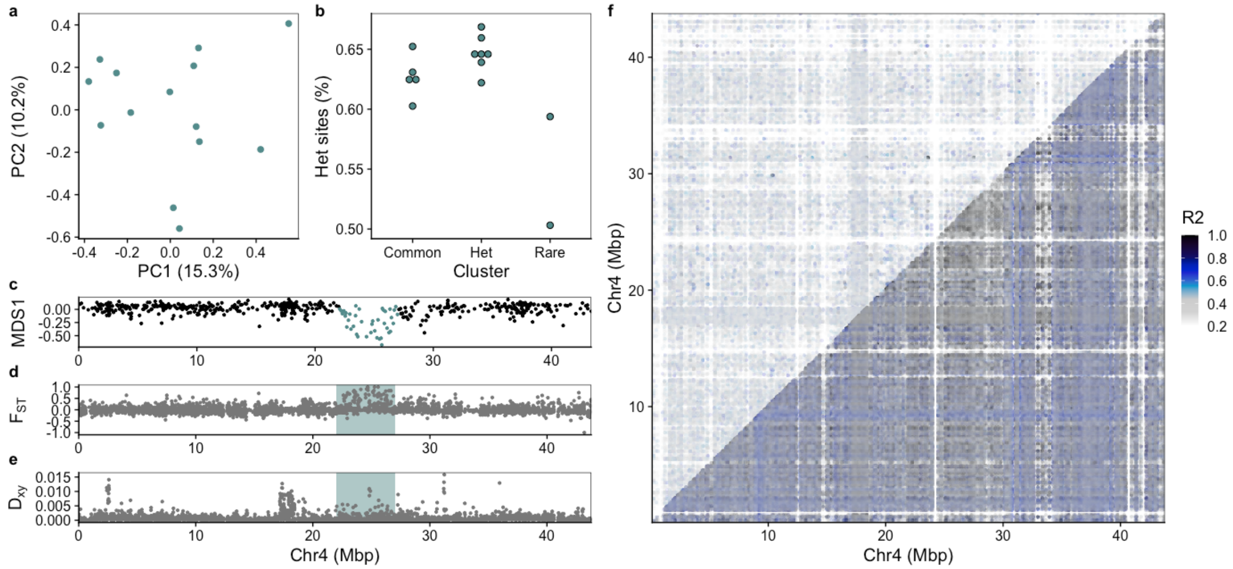


**Supplementary Figure 23:** Putative inversion detected on chromosome 4 (Arctic cod Chr9) using NEAC as reference genome. a) PCA for the inversion region identified using lostruct. b) Manually assigned cluster groups and heterozygous sites given in bins for the clusters. c) MDS analysis produced by lostruct where the inversion region is highlighted. d) $F_{ST}$ and e) $D_{XY}$ calculated with pixy showing elevated values within the highlighted inversion region. f) pairwise linkage disequilibrium plot calculated using pixy where the top triangle includes all samples, and the lower triangle includes only the individuals within the common type.

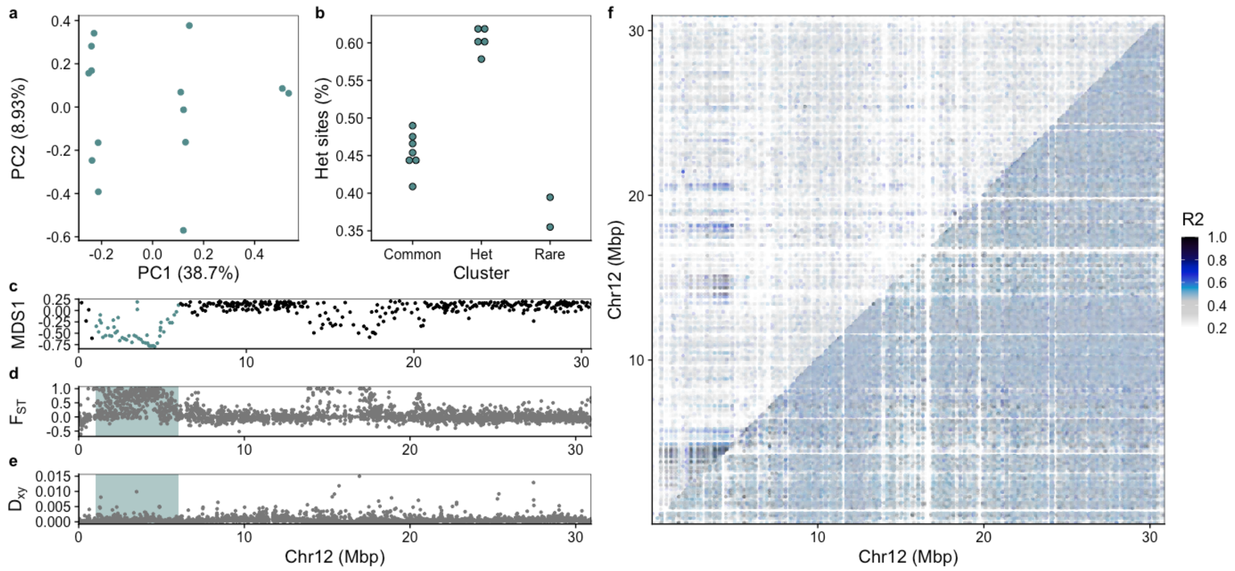


**Supplementary Figure 24:** Inversion detected on chromosome 12 (Arctic cod Chr10) using NEAC as reference genome. a) PCA for the inversion region identified using lostruct. b) Manually assigned cluster groups and heterozygous sites given in bins for the clusters. c) MDS analysis produced by lostruct where the inversion region is highlighted. d) $F_{ST}$ and e) $D_{XY}$ calculated with pixy showing elevated values within the highlighted inversion region. f) pairwise linkage disequilibrium plot calculated using pixy where the top triangle includes all samples, and the lower triangle includes only the individuals within the common type.

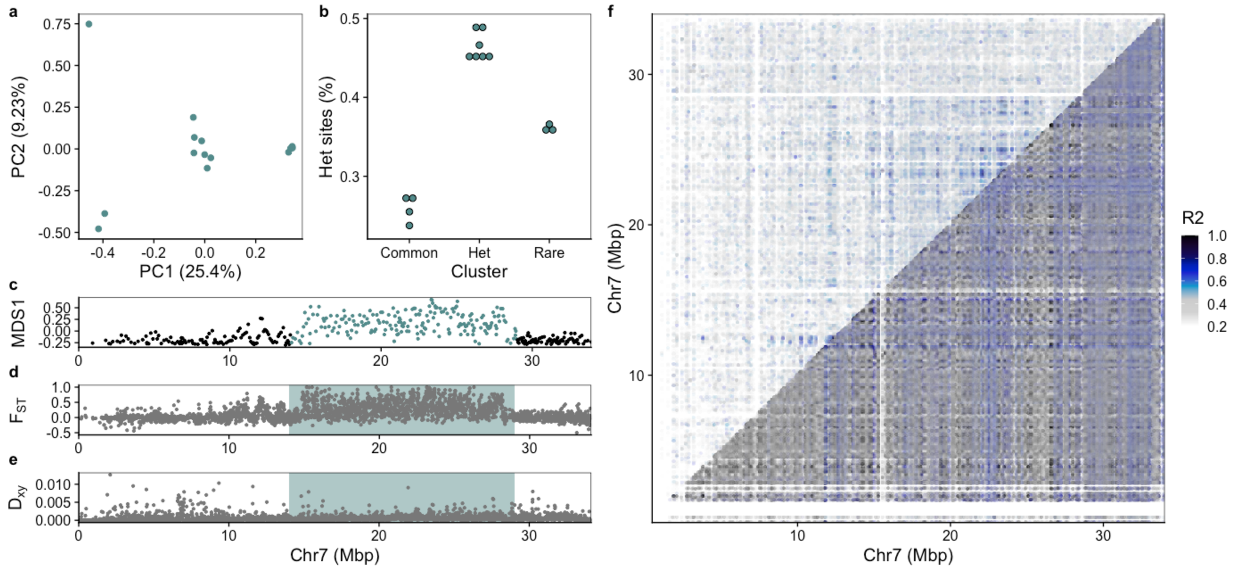


**Supplementary Figure 25:** Inversion detected on chromosome 7 (Arctic cod Chr11) using NEAC as reference genome. a) PCA for the inversion region identified using lostruct. b) Manually assigned cluster groups and heterozygous sites given in bins for the clusters. c) MDS analysis produced by lostruct where the inversion region is highlighted. d) $F_{ST}$ and e) $D_{XY}$ calculated with pixy showing elevated values within the highlighted inversion region. f) pairwise linkage disequilibrium plot calculated using pixy where the top triangle includes all samples, and the lower triangle includes only the individuals within the common type.

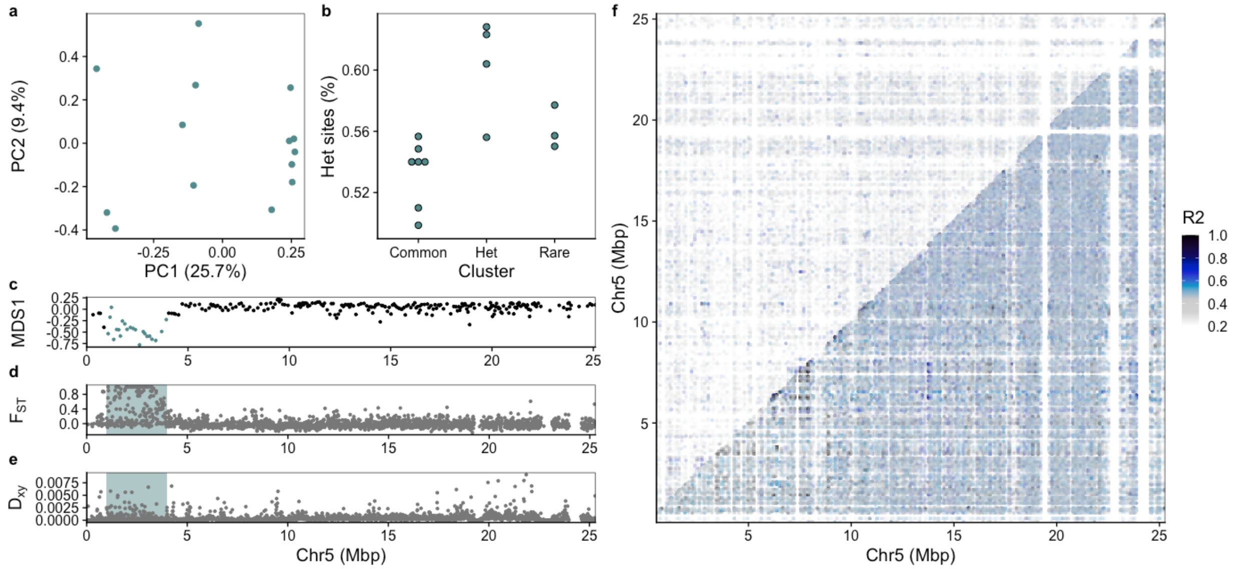


**Supplementary Figure 26:** Inversion detected on chromosome 5 (Arctic cod Chr13) using NEAC as reference genome. a) PCA for the inversion region identified using lostruct. b) Manually assigned cluster groups and heterozygous sites given in bins for the clusters. c) MDS analysis produced by lostruct where the inversion region is highlighted. d) $F_{ST}$ and e) $D_{XY}$ calculated with pixy showing elevated values within the highlighted inversion region. f) pairwise linkage disequilibrium plot calculated using pixy where the top triangle includes all samples, and the lower triangle includes only the individuals within the common type.

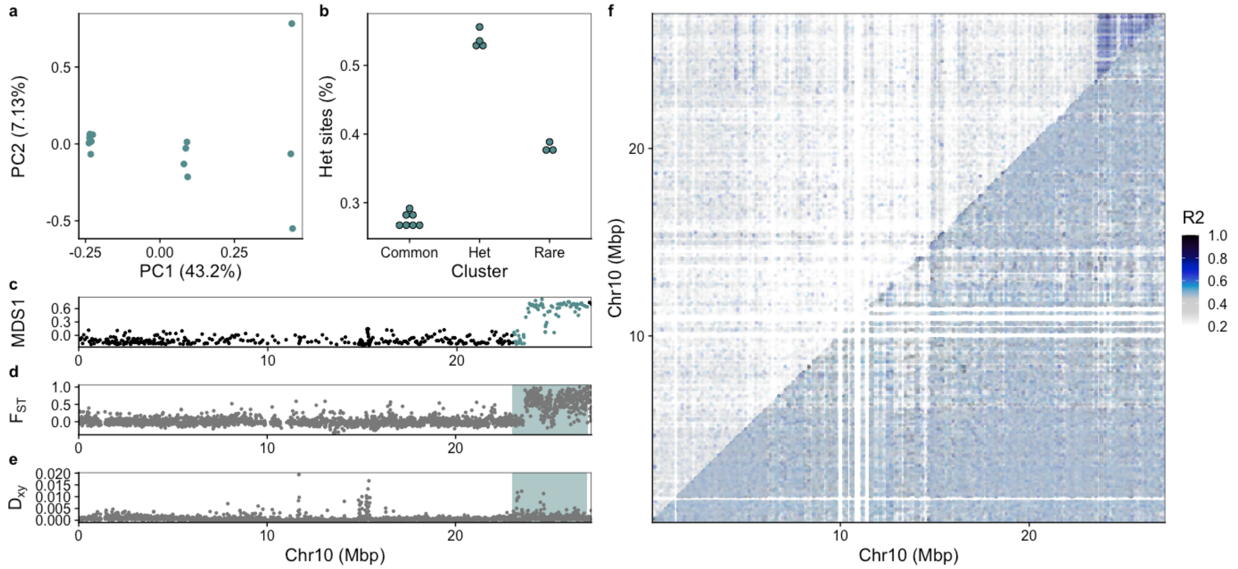


**Supplementary Figure 27:** Inversion detected on chromosome 10 (Arctic cod Chr14) using NEAC as reference genome. a) PCA for the inversion region identified using lostruct. b) Manually assigned cluster groups and heterozygous sites given in bins for the clusters. c) MDS analysis produced by lostruct where the inversion region is highlighted. d) $F_{ST}$ and e) $D_{XY}$ calculated with pixy showing

elevated values within the highlighted inversion region. f) pairwise linkage disequilibrium plot calculated using pixy where the top triangle includes all samples, and the lower triangle includes only the individuals within the common type.

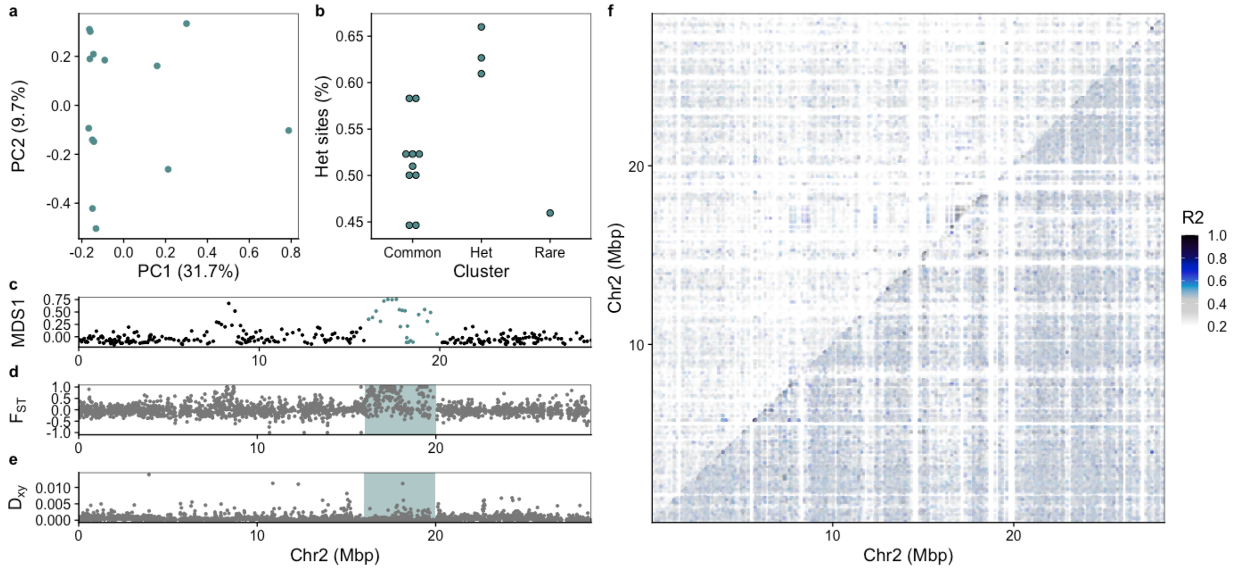


**Supplementary Figure 28:** Putative inversion detected on chromosome 2 (Arctic cod Chr15) using NEAC as reference genome. a) PCA for the inversion region identified using lostruct. b) Manually assigned cluster groups and heterozygous sites given in bins for the clusters. c) MDS analysis produced by lostruct where the inversion region is highlighted. d) $F_{ST}$ and e) $D_{XY}$ calculated with pixy showing elevated values within the highlighted inversion region. f) pairwise linkage disequilibrium plot calculated using pixy where the top triangle includes all samples, and the lower triangle includes only the individuals within the common type.

#### Inversion detection plots using polar cod as reference genome (S29-S41)

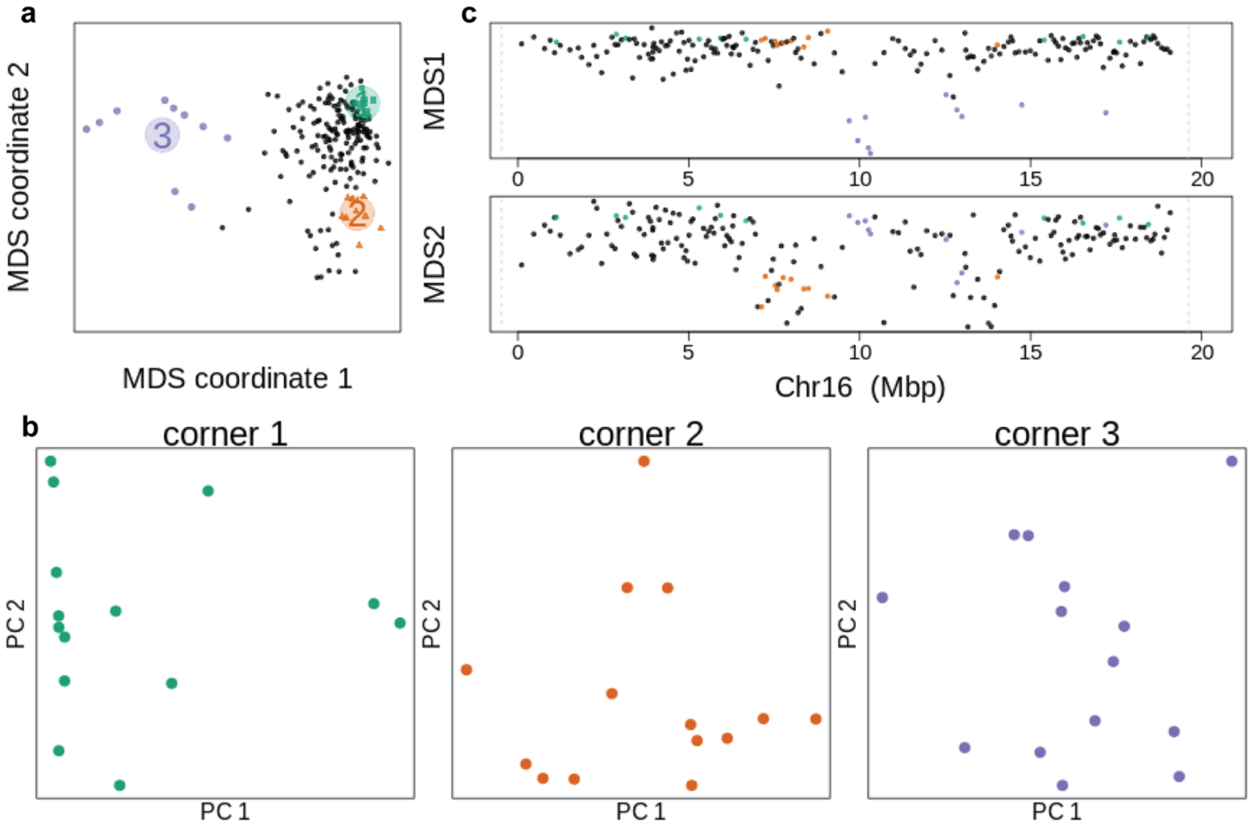


**Supplementary Figure 29:** Lostruct plot showing no putative inversion signal on chromosome 16 (Arctic cod Chr7 (2)) using polar cod as reference genome. a) MDS plots, b) PCA for each MDS corner, and c) PCA windows along the chromosome.

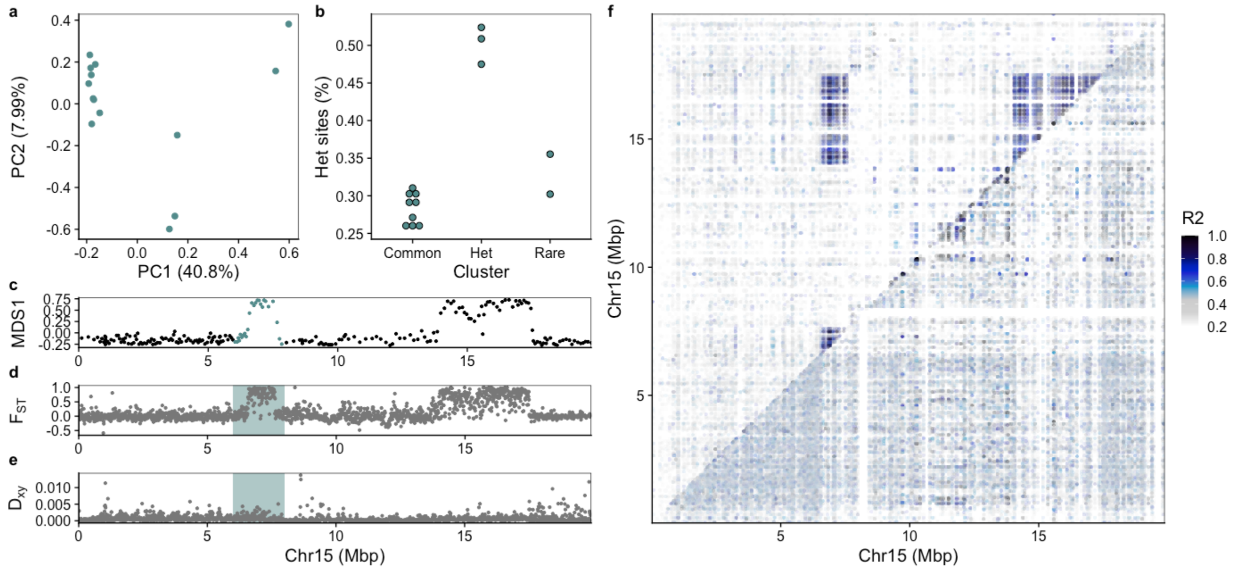


**Supplementary Figure 30:** Split inversion detected on chromosome 15 (Arctic cod Chr1) using polar cod as reference genome. a) PCA for the inversion region identified using lostruct. b) Manually assigned cluster groups and heterozygous sites given in bins for the clusters. c) MDS analysis produced by lostruct where the inversion region is highlighted. d) $F_{ST}$ and e) $D_{XY}$ calculated with pixy showing elevated values within the highlighted inversion region. f) pairwise linkage disequilibrium plot calculated using pixy where the top triangle includes all samples, and the lower triangle includes only the individuals within the common type.

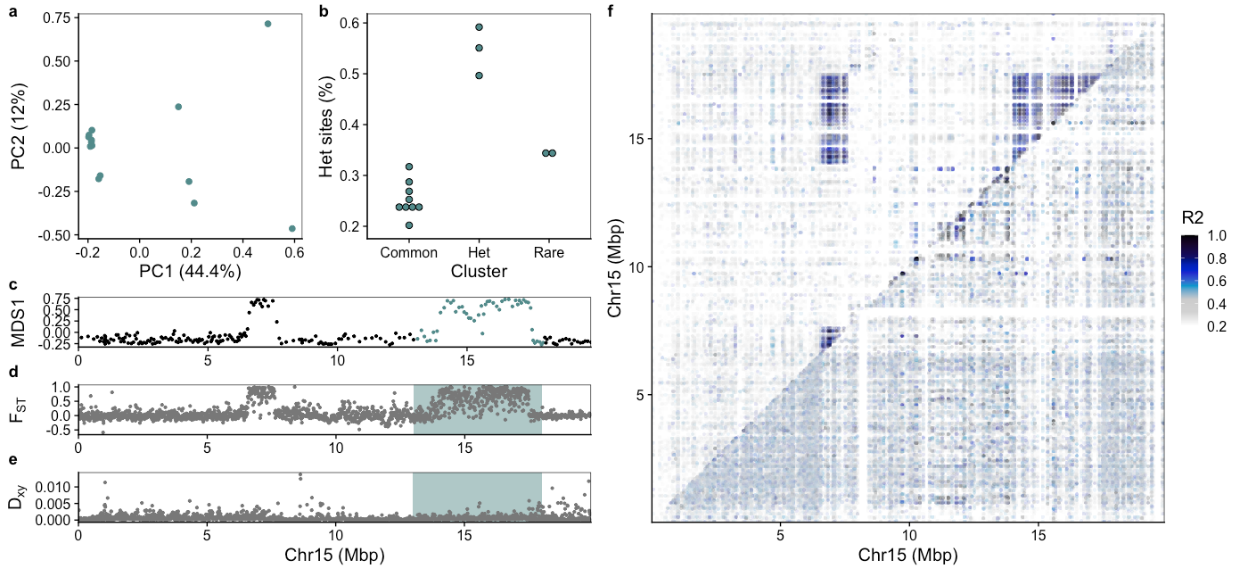


**Supplementary Figure 31:** Split inversion detected on chromosome 15 (Arctic cod Chr1) using polar cod as reference genome. a) PCA for the inversion region identified using lostruct. b) Manually assigned cluster groups and heterozygous sites given in bins for the clusters. c) MDS analysis produced by lostruct where the inversion region is highlighted. d) $F_{ST}$ and e) $D_{XY}$ calculated with pixy showing elevated values within the highlighted inversion region. f) pairwise linkage disequilibrium plot calculated using pixy where the top triangle includes all samples, and the lower triangle includes only the individuals within the common type.

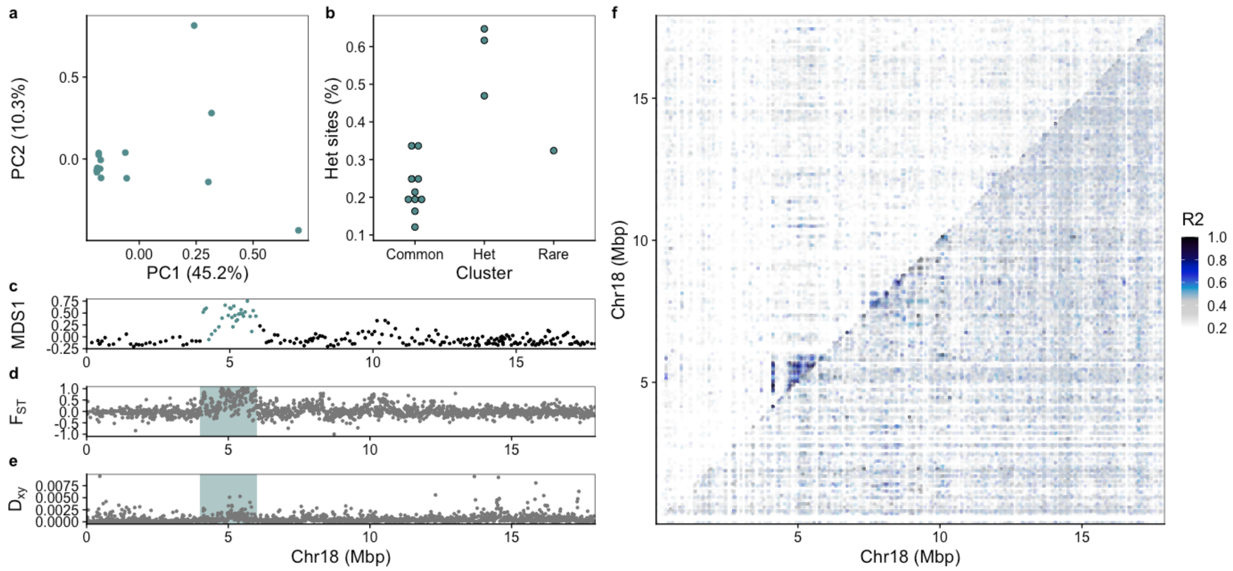


**Supplementary Figure 32:** Putative inversion detected on chromosome 18 (Arctic cod Chr3) using polar cod as reference genome. a) PCA for the inversion region identified using lostruct. b) Manually assigned cluster groups and heterozygous sites given in bins for the clusters. c) MDS analysis produced by lostruct where the inversion region is highlighted. d) $F_{ST}$ and e) $D_{XY}$ calculated with pixy showing elevated values within the highlighted inversion region. f) pairwise linkage disequilibrium plot calculated using pixy where the top triangle includes all samples, and the lower triangle includes only the individuals within the common type.

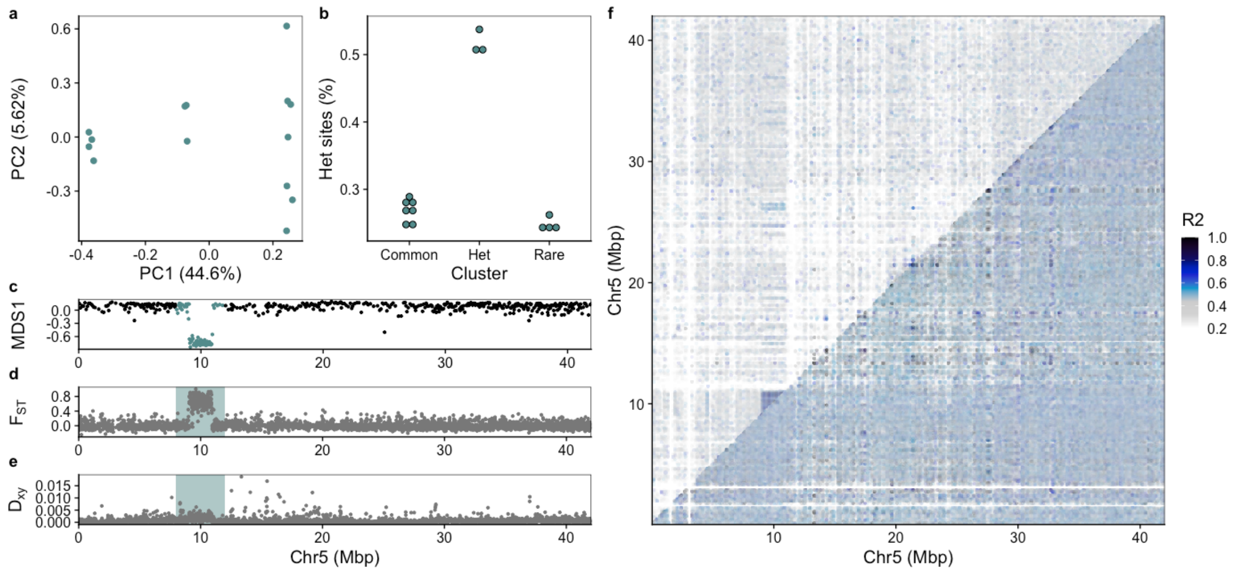


**Supplementary Figure 33:** Inversion detected on chromosome 5 (Arctic cod Chr6) using polar cod as reference genome. a) PCA for the inversion region identified using lostruct. b) Manually assigned cluster groups and heterozygous sites given in bins for the clusters. c) MDS analysis produced by lostruct where the inversion region is highlighted. d) $F_{ST}$ and e) $D_{XY}$ calculated with pixy showing elevated values within the highlighted inversion region. f) pairwise linkage disequilibrium plot calculated using pixy where the top triangle includes all samples, and the lower triangle includes only the individuals within the common type.

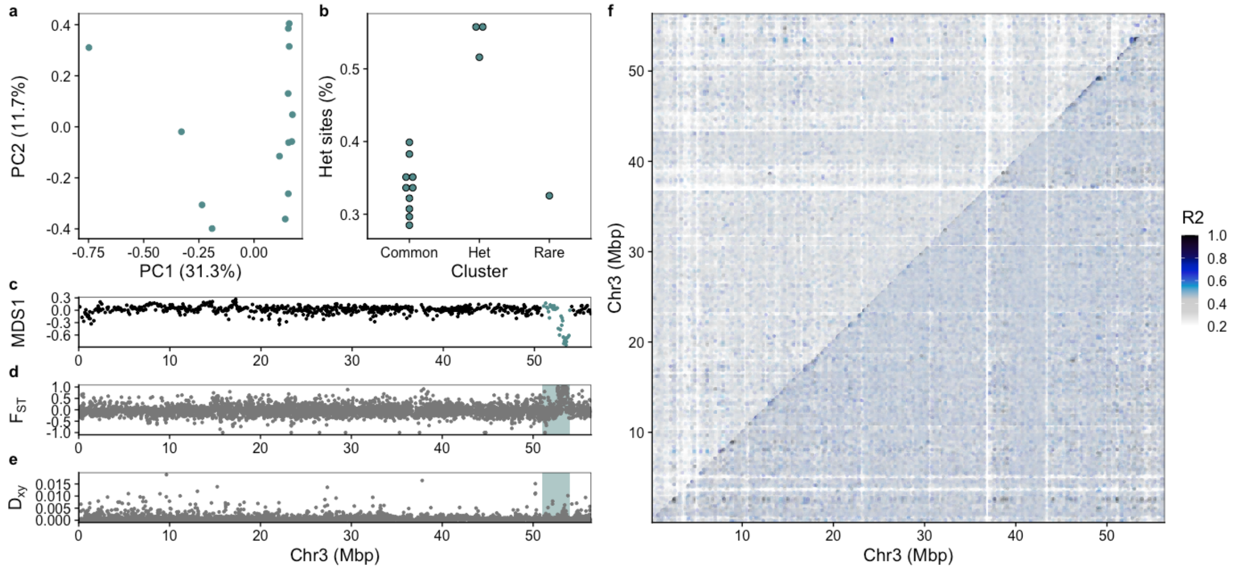


**Supplementary Figure 34:** Putative inversion detected on chromosome 3 (Arctic cod Chr7 (1)) using polar cod as reference genome. a) PCA for the inversion region identified using lostruct. b) Manually assigned cluster groups and heterozygous sites given in bins for the clusters. c) MDS analysis produced by lostruct where the inversion region is highlighted. d) $F_{ST}$ and e) $D_{XY}$ calculated with pixy showing elevated values within the highlighted inversion region. f) pairwise linkage disequilibrium plot calculated using pixy where the top triangle includes all samples, and the lower triangle includes only the individuals within the common type.

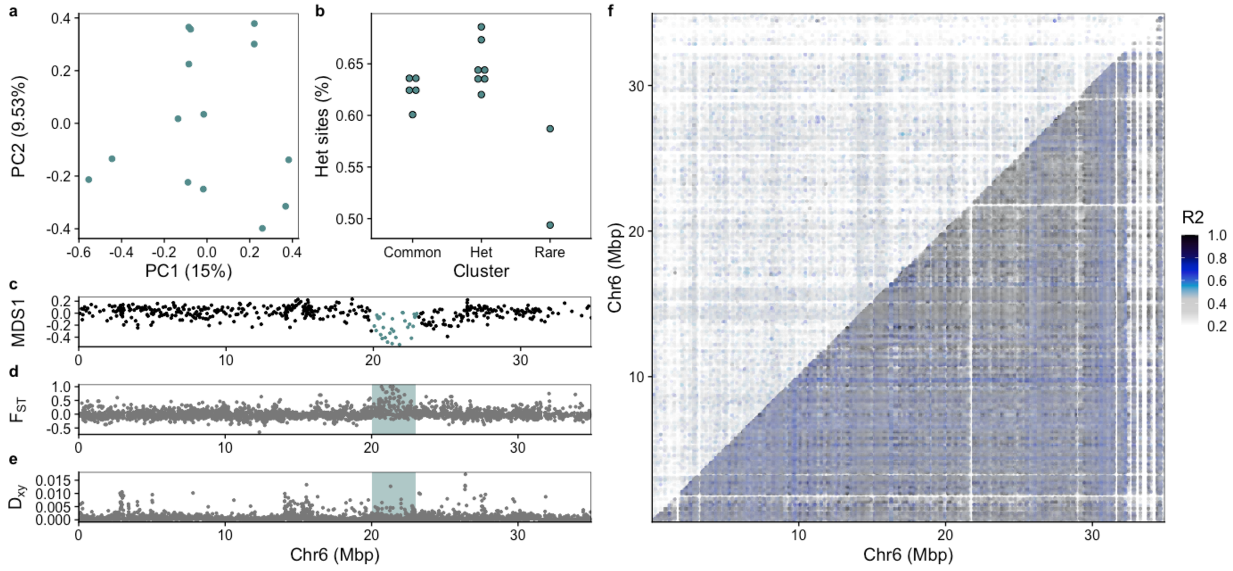


**Supplementary Figure 35:** Putative inversion detected on chromosome 6 (Arctic cod Chr9) using polar cod as reference genome. a) PCA for the inversion region identified using lostruct. b) Manually assigned cluster groups and heterozygous sites given in bins for the clusters. c) MDS analysis produced by lostruct where the inversion region is highlighted. d) $F_{ST}$ and e) $D_{XY}$ calculated with pixy showing elevated values within the highlighted inversion region. f) pairwise linkage disequilibrium plot calculated using pixy where the top triangle includes all samples, and the lower triangle includes only the individuals within the common type.

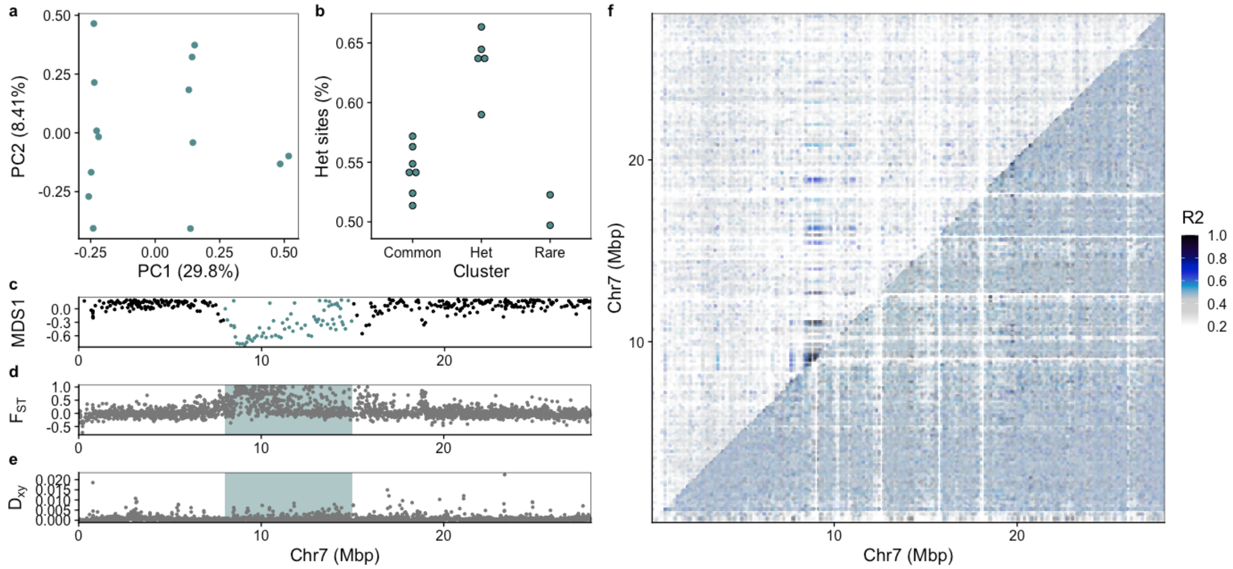


**Supplementary Figure 36:** Inversion detected on chromosome 7 (Arctic cod Chr10) using polar cod as reference genome. a) PCA for the inversion region identified using lostruct. b) Manually assigned cluster groups and heterozygous sites given in bins for the clusters. c) MDS analysis produced by lostruct where the inversion region is highlighted. d) $F_{ST}$ and e) $D_{XY}$ calculated with pixy showing elevated values within the highlighted inversion region. f) pairwise linkage disequilibrium plot calculated using pixy where the top triangle includes all samples, and the lower triangle includes only the individuals within the common type.

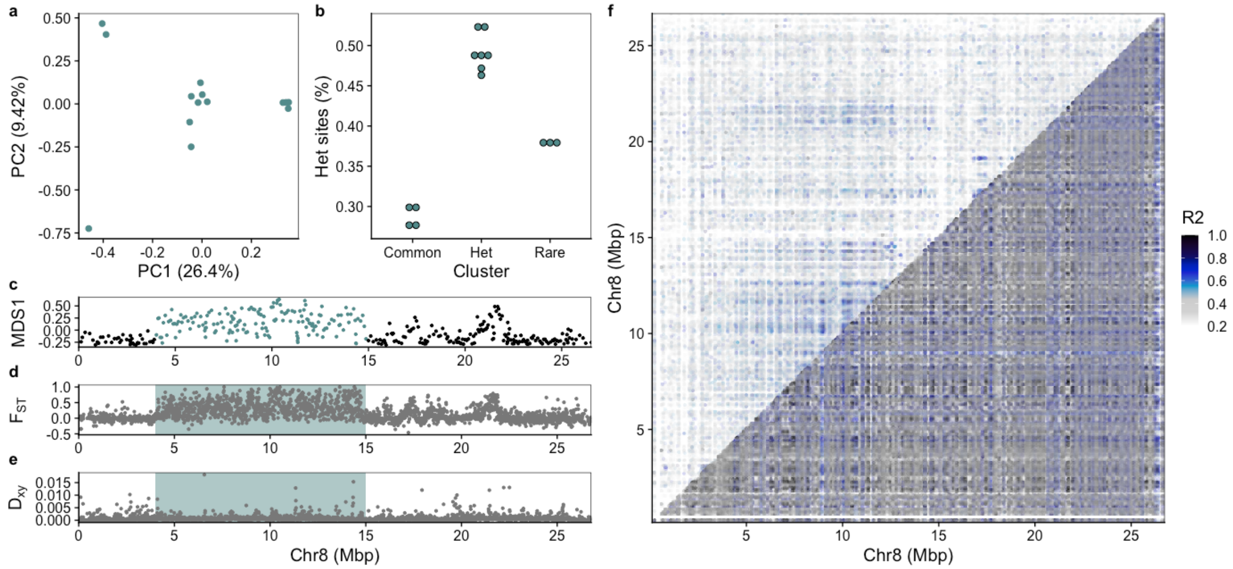


**Supplementary Figure 37:** Inversion detected on chromosome 8 (Arctic cod Chr11) using polar cod as reference genome. a) PCA for the inversion region identified using lostruct. b) Manually assigned cluster groups and heterozygous sites given in bins for the clusters. c) MDS analysis produced by lostruct where the inversion region is highlighted. d) $F_{ST}$ and e) $D_{XY}$ calculated with pixy showing elevated values within the highlighted inversion region. f) pairwise linkage disequilibrium plot calculated using pixy where the top triangle includes all samples, and the lower triangle includes only the individuals within the common type.

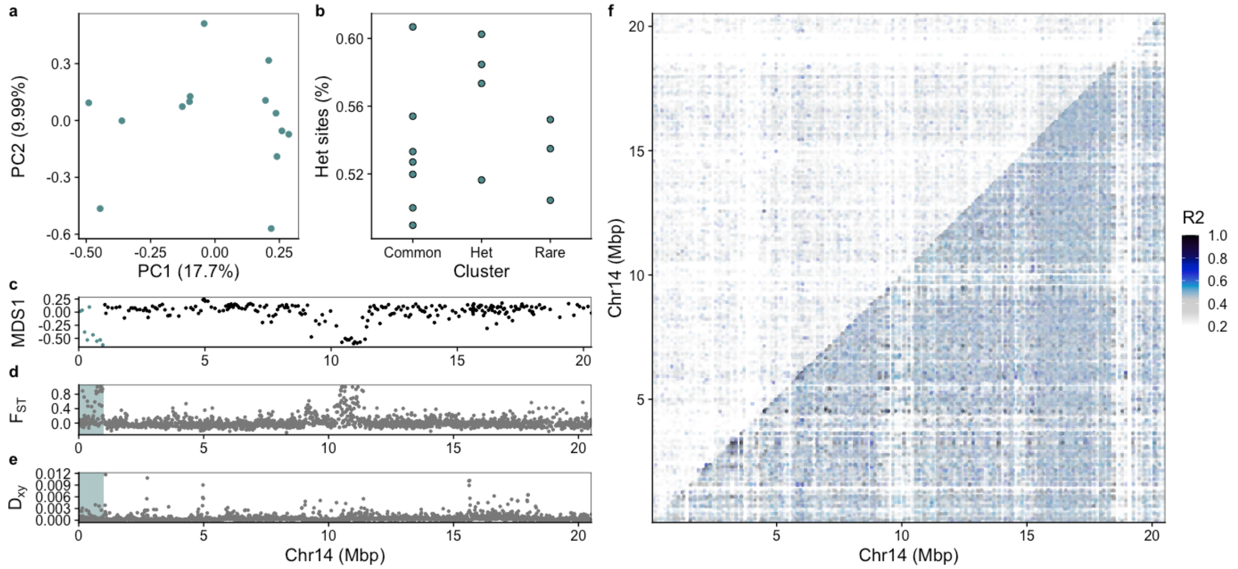


**Supplementary Figure 38:** Split inversion detected on chromosome 14 (Arctic cod Chr13) using polar cod as reference genome. a) PCA for the inversion region identified using lostruct. b) Manually assigned cluster groups and heterozygous sites given in bins for the clusters. c) MDS analysis produced by lostruct where the inversion region is highlighted. d) $F_{ST}$ and e) $D_{XY}$ calculated with pixy showing elevated values within the highlighted inversion region. f) pairwise linkage disequilibrium plot calculated using pixy where the top triangle includes all samples, and the lower triangle includes only the individuals within the common type.

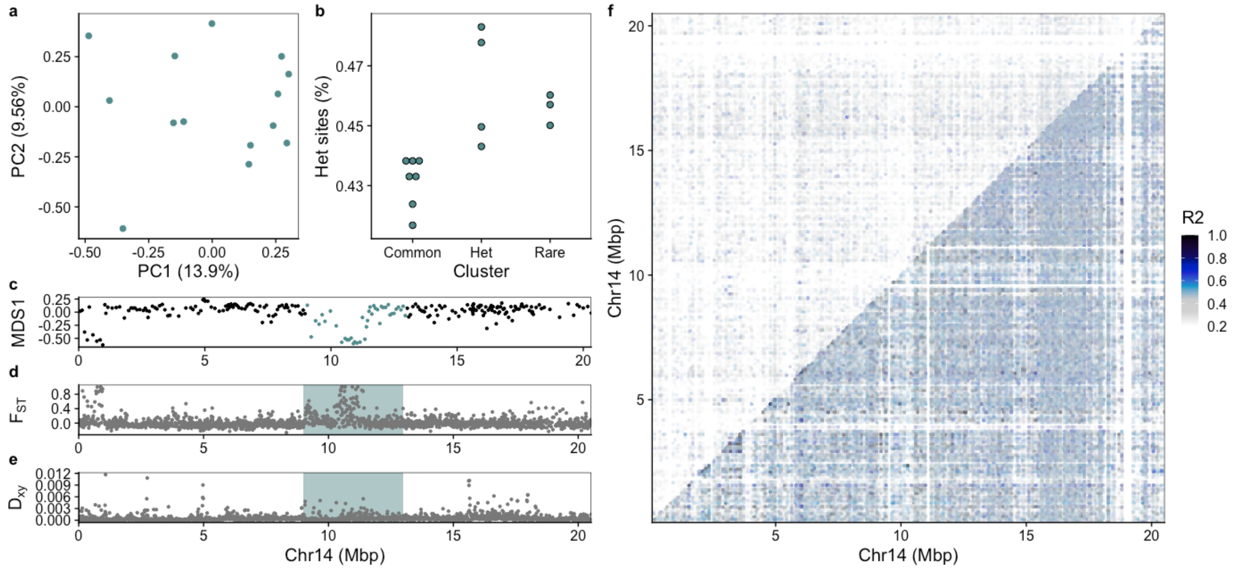


**Supplementary Figure 39:** Split inversion detected on chromosome 14 (Arctic cod Chr13) using polar cod as reference genome. a) PCA for the inversion region identified using lostruct. b) Manually assigned cluster groups and heterozygous sites given in bins for the clusters. c) MDS analysis produced by lostruct where the inversion region is highlighted. d) $F_{ST}$ and e) $D_{XY}$ calculated with pixy showing elevated values within the highlighted inversion region. f) pairwise linkage disequilibrium plot calculated using pixy where the top triangle includes all samples, and the lower triangle includes only the individuals within the common type.

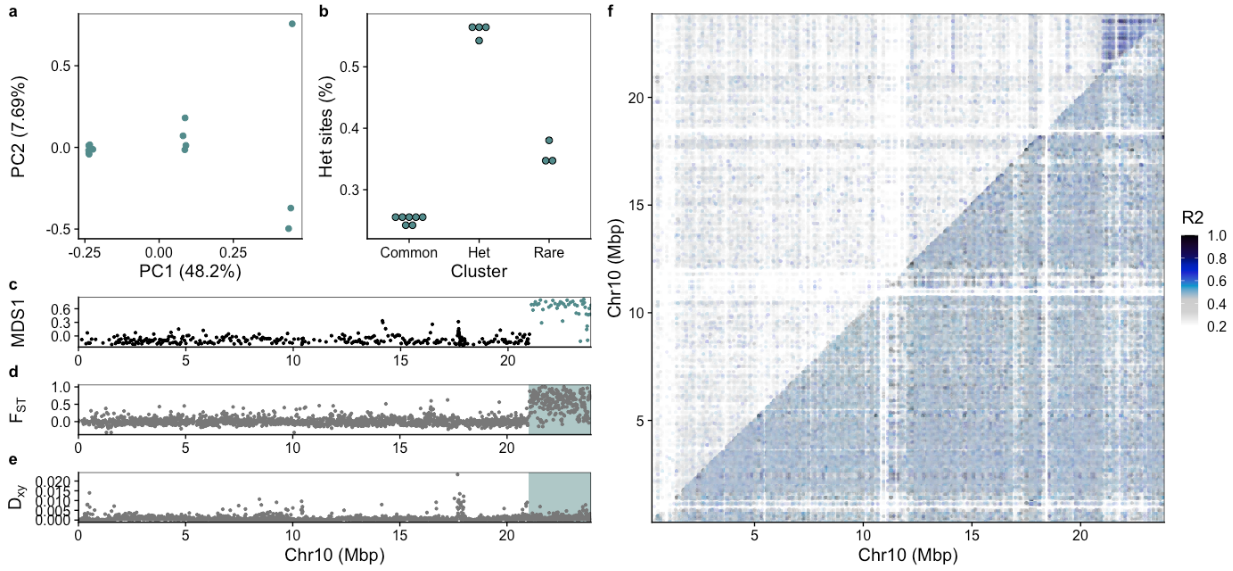


**Supplementary Figure 40:** Inversion detected on chromosome 10 (Arctic cod Chr14) using polar cod as reference genome. a) PCA for the inversion region identified using lostruct. b) Manually assigned cluster groups and heterozygous sites given in bins for the clusters. c) MDS analysis produced by lostruct where the inversion region is highlighted. d) $F_{ST}$ and e) $D_{XY}$ calculated with pixy showing elevated values within the highlighted inversion region. f) pairwise linkage disequilibrium plot calculated using pixy where the top triangle includes all samples, and the lower triangle includes only the individuals within the common type.

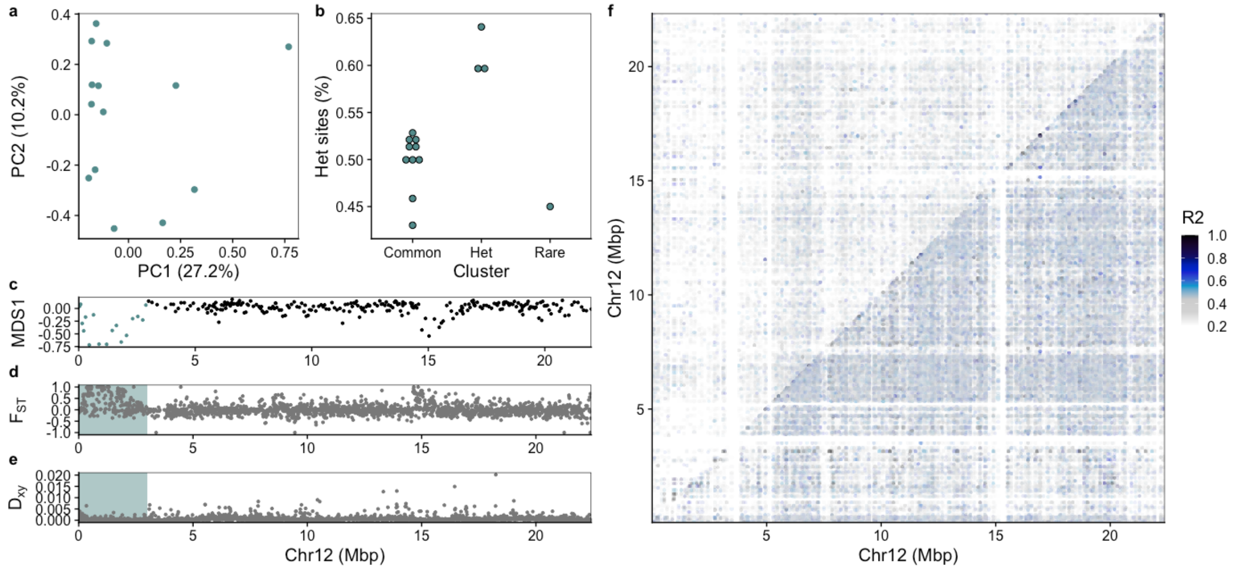


**Supplementary Figure 41:** Putative inversion detected on chromosome 12 (Arctic cod Chr15) using polar cod as reference genome. a) PCA for the inversion region identified using lostruct. b) Manually assigned cluster groups and heterozygous sites given in bins for the clusters. c) MDS analysis produced by lostruct where the inversion region is highlighted. d) $F_{ST}$ and e) $D_{XY}$ calculated with pixy showing elevated values within the highlighted inversion region. f) pairwise linkage disequilibrium plot calculated using pixy where the top triangle includes all samples, and the lower triangle includes only the individuals within the common type.

### Supplementary Tables (T1-T4)

**Supplementary Table 1:** Samples of Arctic cod and polar cod[18] used in this study, including sample ID, length (millimeters), weight (grams), sex, maturity (Mat), trawling locality, DNA extraction protocol, and sample concentration after DNA extraction.

| Sample ID | Length (mm) | Weight (g) | Sex | Mat | Trawling locality | Protocol | Sample concentration (ng/µl) |
|---|---|---|---|---|---|---|---|
| 01-Ag13010 | 206 | 72 | M | 1 | Greenland, Tyroler | Qiagen | 85.05 |
| 02-Ag13014 | 189 | 49.9 | F | 1 | Greenland, Tyroler | Qiagen | 77.13 |
| 03-Ag13019 | 266 | 133 | F | 2 | Greenland, Tyroler | Qiagen | 153.00 |
| 04-Ag13020 | 203 | 56.6 | F | 1 | Greenland, Tyroler | Qiagen | 80.41 |
| 05-Ag13021 | 231 | 82.9 | F | 1 | Greenland, Tyroler | Qiagen | 86.16 |
| 06-Ag13024 | 210 | 67.4 | F | 1 | Greenland, Tyroler | Qiagen | 72.53 |
| 07-Ag13026 | 182 | 45 | F | 0 | Greenland, Tyroler | Qiagen | 69.21 |
| 08-Ag13027 | 204 | 70.2 | M | 1 | Greenland, Tyroler | Qiagen | 64.76 |
| 09-Agl17001 | 203 | 59.55 | F | 1 | Greenland, Besselfjord | Qiagen | 68.43 |
| 10-Agl17002 | 181 | 39.6 | M | 1 | Greenland, Besselfjord | Qiagen | 46.72 |
| 11-Agl17003 | 155 | 21.5 | * | * | Greenland, Besselfjord | Qiagen | 112.46 |
| 12-Ag-ds-0A08045-2 | * | * | * | * | Davis Strait | Qiagen | 7.45 |
| 13-Ag1-hyb-aen-2018 | 131 | 16 | F | 1 | Barents Sea | Qiagen | 239.74 |
| SAMEA4028798+ | * | * | * | * | Davis Strait | ENA: ERR1473882, ERR1473883 | * |
| 04-polcod59 | 15,5 | 24 | F | 2 | Barents Sea | Omega | 14.83 |
| 171-polcod1 | 15 | 22.75 | M | 2 | Barents Sea | Omega | 209.20 |
| 172-polcod2 | 14,1 | 19.35 | M | 2 | Barents Sea | Omega | 138.52 |
| 174-polcod4 | 13,6 | 18.75 | F | 2 | Barents Sea | Omega | 24.15 |
| 178-polcod8 | 15,7 | 24.45 | M | 4 | Barents Sea | Omega | 62.59 |
| 185-polcod20 | 17 | 34 | M | 2 | Barents Sea | Omega | 19.55 |
| 187-polcod37 | 14,3 | 22 | M | 2 | Barents Sea | Omega | 35.96 |
| 188-polcod45 | 17,5 | 35 | F | 3 | Barents Sea | Omega | 16.34 |
| 189-polcod49 | 13,3 | 18 | F | 2 | Barents Sea | Omega | 11.56 |
| 190-polcod50 | 16 | 35 | F | 2 | Barents Sea | Omega | 36.29 |
| 192-polcod52 | 19,5 | 55 | F | 3 | Barents Sea | Omega | 62.70 |
| 228-polcod62 | 18,4 | 49 | M | 4 | Barents Sea | Omega | 131.39 |
| 235-polcod74 | 18,8 | 48 | M | 3 | Barents Sea | Omega | 15.49 |
| 287-polcod31 | 18 | 39 | F | 3 | Barents Sea | Omega | 64.50 |

*No information available. +Accessed from ENA: ERR1473882, ERR1473883.Protocols; Omega: Omega Mag-Bind M6399, Qiagen: QIAGEN Dneasy Blood & Tissue kit.

**Supplementary Table 2:** Filter parameters applied using GATK VariantFiltration. Values indicate at which threshold SNPs were removed.

| VCF | QD | FS | SOR | MQ | MQRankSum | ReadPosRankSum |
|---|---|---|---|---|---|---|
| Intraspecific Arctic cod | < 2 | > 60 | > 3 | < 50 | < -4 | < -5 |
| Cross-species | < 8 | > 60 | > 3 | < 50 | < -10 | < -5 |

**Supplementary Table 3:** Filtering parameters applied using VCFtools. The parameter max missing is on a scale between 0 and 1 where 1 means no missing data is allowed.

| VCF | maf | mac | minQ | max-missing | meanDP (min / max) | DP (min / max) |
|---|---|---|---|---|---|---|
| Intraspecific Arctic cod | 0.075 | 2 | 30 | 0.9 | 8 / 50 | 8 / 50 |
| Cross-species | No filtering | 2 | 30 | 0.9 | 8 / 50 | 8 / 50 |

**Supplementary Table 4:** SNP counts for the different VCFs created in this study.

| VCFs | Number of SNPs | | |
|---|---|---|---|
| | Arctic cod reference | Polar cod reference | NEAC reference |
| Intraspecific Arctic cod | 899623 | 661357 | 721148 |
| Cross-species | 298668 | 275094 | 275248 |

### Supplementary Sequencing Report

DNA samples were processed and sequenced by the Norwegian Sequencing Centre (https://www.sequencing.uio.no). The DNA samples were quantified using the FLUOStar Optima (BMG Labtech) with the Qubit dsDNA HS Assay Kit chemistry (ThermoFisher Scientific). Normalization of all DNA samples to 20ng/ul with Elution Buffer (Qiagen) was performed using the Sciclone G3 NGS Workstation (Perkin Elmer). Normalized DNA samples were sheared using the E220 focused-ultrasonicator (Covaris) with the appropriate manufacturer´s settings for a target fragment mean size of 350bp. After shearing all samples were purified and size selected using KAPA Pure beads (Roche) in a ratio 0.8x (beads:sample) in order to remove fragments shorter than 200bp prior to library preparation.

Library preparation was performed using the KAPA Hyper kit (Roche) on Mosquito LV (Low Volume) pipetting robot (sptlabtech). The library preparation reactions (End repair, A-tailing and adapter Ligation) were performed using 5x reduced volume compared to the kit reaction volumes. The IDT for Illumina TruSeq DNA UD 96 Indexes (Illumina) were used for barcoding each 96-plate of samples. After the ligation of adapters, the samples volume was increased to 25ul with the addtion of EB buffer (Qiagen), and one round of bead cleanup with ratio 0.8x was performed. The libraries were subsequently amplified with 5 cycles of PCR. The PCR reactions were done in 2x reduced volume compared to the kit PCR reactions. All incubations were executed according to the manufacturer's instructions. The final libraries were purified, and size selected using KAPA Pure beads (Roche) in a ratio 0.8x.

After library preparation and cleanup, all libraries were run on a 5200 Fragment Analyzer System (Agilent) using the NGS Fragment Kit: DNF-473-0500 (Agilent) for determination of the average size of each library. Subsequently, absolute quantification of each library was done using the KAPA Library Quanitification Kits (Roche), on a LightCycler 480 qPCR instrument (Roche) in 10ul reaction volume. Finally, after determining the absolute concentration of each library (in nM) using the Fragment analyzer and qPCR results, all libraries were normalized to the same molarity using the Sciclone G3 NGS Workstation (Perkin Elmer) and equal volumes of each sample were pooled, creating 96plex pools. Each of the pools was sequenced on several lanes of a HiSeq4000 System (Illumina) in 2x150bp mode (150bp Paired End), using a HiSeq 3000/4000 SBS Kit (300 cycles) (Illumina).
